# CDX4 regulates the progression of neural maturation in the spinal cord

**DOI:** 10.1101/177469

**Authors:** Piyush Joshi, Andrew J. Darr, Isaac Skromne

## Abstract

The progressive maturation of cells down differentiation lineages is controlled by collaborative interactions between networks of extracellular signals and intracellular transcription factors. In the vertebrate spinal cord, FGF, Wnt and Retinoic Acid signaling pathways regulate the progressive caudal-to-rostral maturation of neural progenitors by regulating a poorly understood gene regulatory network of transcription factors. We have mapped out this gene regulatory network in the chicken pre-neural tube, identifying CDX4 as a dual-function core component that simultaneously regulates gradual loss of cell potency and acquisition of differentiation states: in a caudal-to-rostral direction, CDX4 represses the early neural differentiation marker *Nkx1.2* and promotes the late neural differentiation marker *Pax6.* Significantly, CDX4 prevents premature PAX6-dependent neural differentiation by blocking *Ngn2* activation. This regulation of CDX4 over *Pax6* is restricted to the rostral pre-neural tube by Retinoic Acid signaling. Together, our results show that in the spinal cord, CDX4 is part of the gene regulatory network controlling the sequential and progressive transition of states from high to low potency during neural progenitor maturation. Given CDX well-known involvement in *Hox* gene regulation, we propose that CDX factors coordinate the maturation and axial specification of neural progenitor cells during spinal cord development.

## INTRODUCTION

Differentiating cells transition from one temporary state to another, losing potency and acquiring specialized functions in the process. Each step along the differentiation pathway is defined by a unique assortment of active transcription factors (Davidson, 2006; Royo et al., 2011). This transcriptome can change over time, mostly cued by dynamic extra-cellular signaling factors (Peter and Davidson, 2013; Sandmann et al., 2007). It is the cross-regulation between transcription and signaling components that promotes the progressive acquisition of specialized functions while preventing dedifferentiation: transcription factors specify the cell’s identity and ability to respond to signaling factors (competence), and signaling factors control the sequential activity of transcription factors to promote directional acquisition of specialized traits (Davidson and Levine, 2008; Levine and Davidson, 2005; Sandmann et al., 2007). These interactions between transcription factors and signaling pathways form complex networks that have been challenging to dissect, hindering our understanding of the mechanisms regulating cellular state transitions.

The vertebrate spinal cord serves as an important accessible model to study the maturation of neural progenitors during their transition from one cellular state to the next. Maturation of spinal cord progenitors at the caudal end of the embryo follows a caudal-to-rostral organization, with undifferentiated cells localizing to the caudal regions and more mature cells localizing to the more rostral positions (Butler and Bronner, 2015; Diez del Corral et al., 2003; Diez del Corral and Storey, 2004; Wilson et al., 2009). During the early segmentation stages in chick embryos up to the point of tailbud formation (0 to 16 somites equivalent to Hamburger and Hamilton (HH) stages 6-12; Hamburger and Hamilton, 1951), extensive fate mapping and gene expression analysis has resulted in the identification of four distinct embryonic regions corresponding to four different neural maturation states (reviewed in Gouti et al., 2015 and Henrique et al., 2015; summarized in Fig 1A). The most caudal region is the caudal lateral epiblast and node-streak border region containing bipotent neuromesodermal progenitors (NMPs) cells that contribute to both neural and mesodermal tissues (region 1; Brown and Storey, 2000; Cambray and Wilson, 2007; Tzouanacou et al., 2009; reviewed in Henrique et al., 2015). NMPs are defined molecularly by the co-expression of two key transcription factors, the pan-neural marker *Sox2* and the mesodermal marker *T/Bra,* although NMPs also transcribe the pre-neural identity marker *Nkx¡.2* (also known as *Sax1;* Delfino-Machin et al., 2005; Gouti et al., 2015; Gouti et al., 2017). Immediately rostral to the NMP domain is the pre-neural tube (PNT; Gouti et al., 2015; Henrique et al., 2015). Cells in the PNT downregulate *T/Bra* but continue to express *Sox2* (Delfino-Machin et al., 2005; Gouti et al., 2015). PNT can be further subdivided into a caudal PNT that continues to express *Nkx1.2* (region 2) and the rostral PNT which downregulates *Nkx1.2* and activates *Pax6* transcription (region 3; Bel-Vialar et al., 2007; Bertrand et al., 2000; Delfino-Machin et al., 2005; Sasai et al., 2014; Spann et al., 1994). Finally, rostral to the PNT and situated adjacent to the developing somites is the neural tube (NT; region 4; Gouti et al., 2015; Henrique et al., 2015). NT cells are *Nkx1.2*’-negative and Pax6-positive and begin to transcribe the neural differentiation genes *Ngn1/2* and *NeuroM*(Diez del Corral et al., 2003). Thus, from caudal-to-rostral, four spatially distinct populations can be identified that correspond to four maturation states (summarized in Fig 1A).

**Fig 1.**
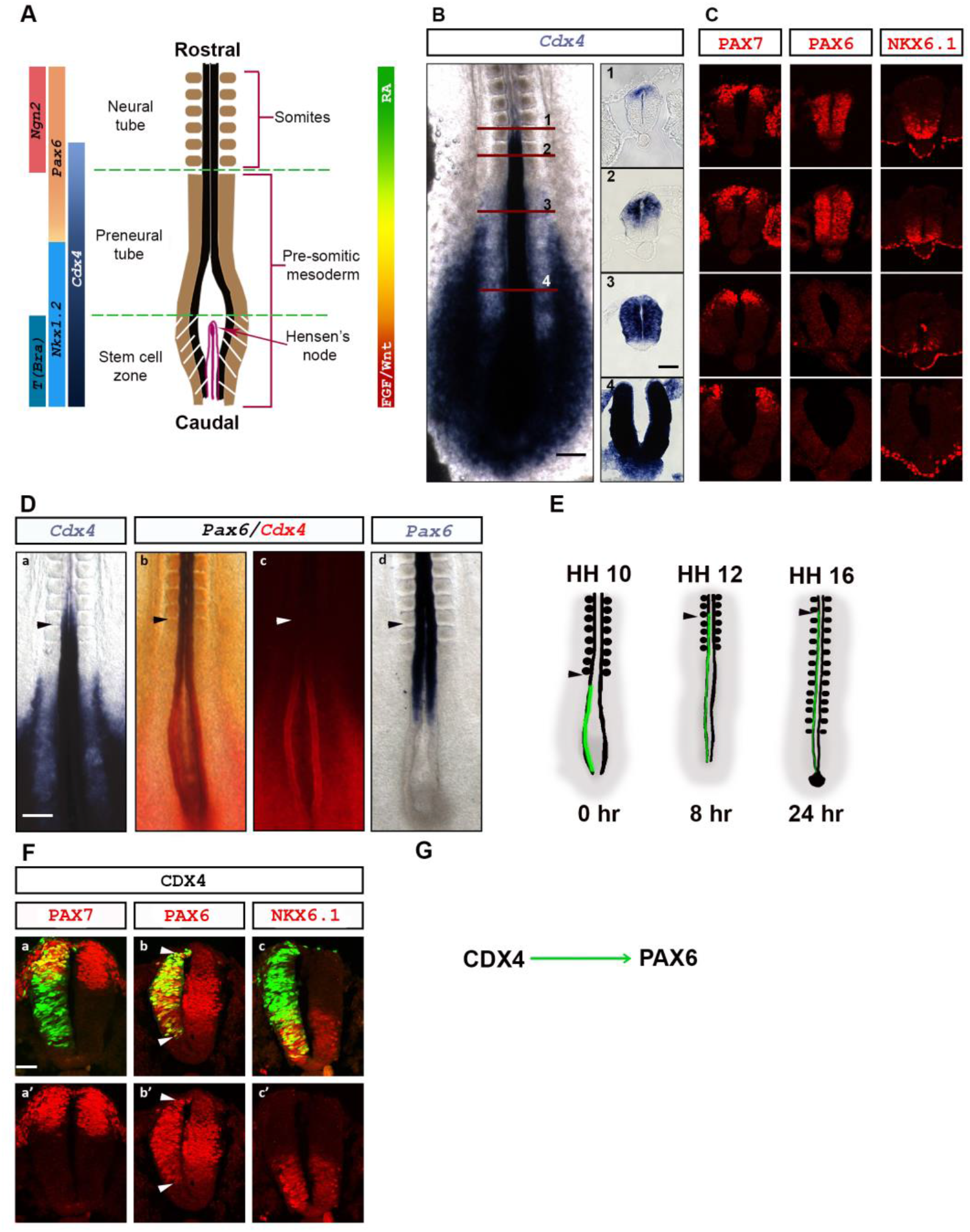
CDX4 activates transcription of the neurogenesis gene *Pax6.* (**A**) Schematic representation of the caudal end of HH10 chicken embryo showing primary subdivisions (central diagram; adapted from Olivera-Martinez and Storey, 2007), and expression domains of key transcription and signaling factors (left and right of diagram, respectively). (**B**) *Cdx4* is transcribed in a dynamic dorsal-ventral (DV) gradient along the rostro-caudal (RC) axis of embryos (HH11). Red lines indicate position of transverse sections shown on right. (**C**) Distribution of PAX7 (dorsal), PAX6 (dorsal-to-intermediate), and NKX6.1 (ventral) proteins relative to *Cdx4* transcription domain. (**D**) *Cdx4* and *Pax6* transcription domains overlap in the rostral pre-neural tube (PNT) at stage HH 11-11+ (ISH; *Cdx4* expression in purple in Da, and red in Db and Dc; *Pax6* expression in purple in Db and Dd). Arrow shows position of somite 13. (**E**) Graphical representation of the experimental approach used throughout this work. The PNT of HH10-11 stage embryos were electroporated on the left side with appropriate constructs carrying a GFP-reporter gene. Embryos were processed for analysis 8 (HH 12-13) or 24 (HH16-16+) hours post electroporation (hpe), when electroporated cells are localized to the rostral portion of the PNT (HH 12-13) or the caudal portion of the NT (thoracic level; HH16-16+), respectively. Control experiments demonstrate that electroporation alone has not affect gene transcription, and that overexpression of electroporated constructs is long lasting (Fig S1 for all experiments). Arrowhead shows the position the last somite formed at the time of electroporation. (**F**) CDX4 does not regulate DV patterning in the neural tube. Ectopic *Cdx4* did not change the distribution of PAX7 or NKX6.1 proteins (n=6/6 for both), but caused ectopic PAX6 accumulation outside its normal domain (arrowheads, n=6/6). Marker proteins are in red and electroporated cells are in green (nuclear GFP tag). Embryos were electroporated at HH10-11 and analyzed 24 hpe (HH16). (E) Summary of results. Scale bar is 200μm for whole mount and 40μm for transverse sections.

Recently, single cell transcriptome analysis of *in vitro* differentiating NMPs has confirmed and expanded the known transcriptional signatures observed throughout NMPs initial cell fate choice decision, allowing the more accurate assignment of gene activities to particular specification states (Gouti et al., 2014; Gouti et al., 2017). For example, CDX transcription factors have been implicated in NMP maintenance, and also axial patterning and elongation (Amin et al., 2016), however the separation of these activities has been challenging to dissect due to the multiple and partially redundant activities of the three functionally similar CDX proteins (CDX1, CDX2 and CDX4; van Rooijen et al., 2012). By studying the differentiation of NMPs derived from mouse Embryonic Stem Cells (mESC) lacking individual or different combination of *Cdx* genes, CDX proteins were shown to regulate the temporal maintenance of *T/Bra* (Gouti et al., 2017). By maintaining or down regulating *T/Bra* transcription, *Cdx* genes regulate the fate decision of NMP cell to become either mesoderm or neural tissues (Gouti et al., 2017). In addition to the NMPs, *Cdx* are also transcribed in NMP descendants in the PNT and NT (Gaunt et al., 2005; Marom et al., 1997; Fig 1B), where their function remains largely unknown.

The transition from NMP to pre-neural to neural transcriptional states is under the control of three signaling factors: FGF, Wnt and Retinoic Acid (RA; Diez del Corral et al., 2003; Olivera-Martinez and Storey, 2007). At the caudal end of the embryo, FGF8 and Wnts (Wnt3a and Wnt8c) are transcribed in a caudal to rostral gradient that promotes potency by maintaining *T/Bra, Sox2* and *Nkx1.2* expression while simultaneously preventing *Pax6* transcription (Bertrand et al., 2000; Delfino-Machin et al., 2005; Diez del Corral et al., 2003; Olivera-Martinez et al., 2012). FGF also maintains tissue proliferation by limiting precocious cell cycle exit (Akai et al., 2005). In contrast, RA secreted from somites establishes a rostral to caudal signaling gradient that promotes differentiation: first by promoting transcription of neural identity genes *Pax6* (Diez del Corral et al., 2003; Novitch et al., 2001; Pituello et al., 1999), and subsequently, by promoting transcription of downstream neurogenic genes *Ngn1/2* and *NeuroM* (Diez del Corral et al., 2003). By inducing *Pax6* and *Ngn2,* RA induces cells to exit the proliferation program (Bel-Vialar et al., 2007; Lacomme et al., 2012). The signaling activities of FGF/Wnt and RA are segregated to opposite caudal and rostral regions of the nascent spinal cord through positive and negative interactions: caudally, high FGF directly prevents RA synthesis and stimulates its degradation, while rostrally, low FGF indirectly promotes RA production through a Wnt8c-dependent mechanism (Boulet and Capecchi, 2012; Olivera-Martinez et al., 2012; Olivera-Martinez and Storey, 2007; Sakai et al., 2001; White et al., 2007). In turn, RA inhibits *Fgf8* transcription rostrally, creating a zone where cells can exit the cell cycle and differentiate (Diez del Corral et al., 2003; Kumar and Duester, 2014). These interactions have been proposed to function as the signaling switch that drives the transition of cellular states in the caudal neural tube (Diez del Corral and Storey, 2004; Olivera-Martinez and Storey, 2007). While the signal interactions regulating the transition from NMP to pre-neural to neural states have been extensively investigated, the underlying transcription factor network driving the cell transitions are incompletely understood.

FGF, Wnt and RA signals are known regulators of *Cdx* transcription (Deschamps and van Nes, 2005; Lohnes, 2003), making CDX transcription factors good candidates to regulate PNT cell maturation. In chicken embryos, *Cdx* genes are transcribed in nested domains, at levels that are high in NMPs and low in the NT (Marom et al., 1997). These high-to-low levels of transcription have also been observed in differentiating NMPs *in vitro* (Gouti et al., 2017). In the spinal cord, CDX factors are essential for tissue specification and rostro-caudal patterning (Deschamps et al., 1999; Nordstrom et al., 2006; Shimizu et al., 2006; Skromne et al., 2007; van den Akker et al., 2002), controlling the initial specification of post-occipital tissues (van Rooijen et al., 2012), and the subsequent patterned transcription of *Hox* expression domains (Deschamps et al., 1999; Hayward et al., 2015). Thus, *Cdx* genes are attractive candidates to integrate multiple signals into coherent cell maturation states.

Here we show that chicken CDX4, the only CDX present in the chick PNT and NT during early segmentation stages (HH10-16; Marom et al., 1997), controls the progression of PNT cells towards more mature states without promoting their terminal differentiation. In the PNT, transient CDX4 results in *Nkx1.2* downregulation and *Pax6* activation, which drives cells with recently acquired neural identity (*Sox2+, Nkx1.2+, Pax6-*) toward a more restricted neural progenitor state (*Sox2+, Nkx1.2-, Pax6+).* Significantly, *Pax6* activation by CDX4 is dependent on RA secreted by somites, which restricts the maturation of cells to the rostral PNT. Furthermore, we show that CDX4 prevents *Ngn2* transcription even in the presence of the *Ngn2-* activator PAX6, thus preventing the premature cell’s terminal differentiation. Our results support a model in which CDX4 is an integral component of a gene regulatory network that functions to simultaneously reduce the potency and increase the differentiation state of cells. We propose that this gene regulatory network operates under the control of previously described signaling network involving FGF, Wnt, and RA.

## RESULTS

### *Cdx4* is transcribed in the caudal neural tube where it regulates *Pax6* transcription

CDX4 neural function in chicken embryos was first analyzed by correlating its transcription domain to distinct progenitor cell maturation zones of the caudal neuroectoderm (Fig 1A; Olivera-Martinez and Storey, 2007). As previously reported in whole chick embryos (HH10-12; Morales et al., 1996; Marom et al., 1997), *Cdx4* is transcribed in the pre-neural tube (PNT) and nascent neural tube (NT) in a high caudal to low rostral gradient (Fig 1B). However, transverse sections also revealed that *Cdx4* is transcribed in a highly dynamic dorsal-to-ventral (DV) gradient: caudally, *Cdx4* transcription was ubiquitous throughout the medio-lateral extent of the PNT (dorsal-ventral extent in the NT), whereas rostrally, *Cdx4* transcription was progressively excluded from ventral regions as well as the roof plate (Fig 1B, transverse sections). Due to the lack of chicken specific CDX4 antibody, we were unable to examine the CDX4 protein profile in the neural tube. However, a similar dorsally restricted expression profile has been reported for *Cdx4* in mouse embryos (Gaunt et al., 2005), suggesting evolutionary conserved transcriptional mechanisms and a potential function for CDX4 in the specification of DV neural cell identities.

To test the role of CDX4 in DV specification, we analyzed *Cdx4* transcriptional domain relative to various DV identity markers including the dorsal cell marker *Pax7* (Briscoe et al., 2000; Diez del Corral et al., 2003), the dorsal-to-intermediate cell marker *Pax6* (Briscoe et al., 2000; Novitch et al., 2003), and the ventral cell marker *Nkx6.1* (Briscoe et al., 2000; Diez del Corral et al., 2003; Novitch et al., 2003). At HH11, we observed a correlation between the transcriptional domain of *Cdx4* and some of these markers. For example, in the caudal NT, PAX7 domain was nested within, and NKX6.1 domain was complementary to *Cdx4* domain of transcription (Fig 1B, C). However, more rostrally in the NT, we saw a loss of correlation between D/V markers and *Cdx4* domain of transcription; PAX7 domain was broader than, and NKX6.1 domain no longer complemented *Cdx4* transcription domain (Fig 1B, C). The only correlation we observed was between *Cdx4* and *Pax6,* with levels of *Cdx4* transcript decaying as levels of *Pax6* transcript and PAX6 protein increased in a caudal to rostral direction (Fig 1C, D).

To formally test *Cdx4* involvement in DV cell fate specification, we artificially maintained high levels of *Cdx4* in the NT in a domain where *Cdx4* would normally be downregulated. We reasoned that if CDX4 regulates DV cell specification, increasing *Cdx4* levels would result in a change in the localization of DV marker genes. We overexpressed CDX4 by electroporating wild type *Cdx4* in the PNT of stage HH10-11 embryos, a region that transcribes endogenous *Cdx4,* and analyzed the protein distribution of PAX7, PAX6, and NKX6.1 24-hours post-electroporation (hpe; HH16-17), at a time when electroporated cells have become part of the NT and no longer transcribe endogenous *Cdx4* (Fig 1B, E). While artificially maintained high levels of *Cdx4* expression did not change NKX6.1 and PAX7 protein distribution (Fig 1F; n=6/6 for both conditions), ectopic *Cdx4* caused production of PAX6 protein outside its normal domain, both ventrally and dorsally (Fig 1F; n=6/6). In this and all other experiments, electroporation of a control reporter GFP vector had no effect on target gene transcription and protein distribution (Fig S1). Together, these results suggest that CDX4 is not a general regulator of DV identity markers, but instead, a specific regulator of *Pax6* transcription (Fig 1G).

### CDX4 regulates *Pax6* transcription during neural progenitor cell maturation

In addition to its function in DV cell specification, PAX6 promotes the maturation of neural progenitor cells in the PNT (Bel-Vialar et al., 2007). Given that our results do not support a function for CDX4 in global DV cell specification (Fig 1), we hypothesized that CDX4 might regulate *Pax6* transcription during PNT cell maturation. To test this hypothesis, we asked whether the presence of CDX4 was sufficient to change *Pax6* transcription in the rostral PNT, a region where *Pax6* transcription initiates. Embryos were electroporated in the PNT with different *Cdx4* constructs (HH10-11), grown for 8 hours only (HH12-13), and analyzed by *in situ* hybridization for premature *Pax6* activation. Two constructs were used in this assay, a wild type and a constitutive active version of CDX4 that phenocopies CDX functions in *Hox* gene transcription assays (VP16CDX4; Bel-Vialar et al., 2002; Faas and Isaacs, 2009). In these short incubation experiments, VP16CDX4 was able to induce *Pax6* transcription more caudally and at higher levels than CDX4 (Fig 2A; n=4/6 by ISH. Fig 2B; n=3/4 by IHC), while CDX4 could induce *Pax6* after long incubation periods (24 hpe; Fig 1F). These results suggest that CDX4 has the potential to regulate *Pax6* transcription in the rostral PNT and caudal NT.

**Fig 2.**
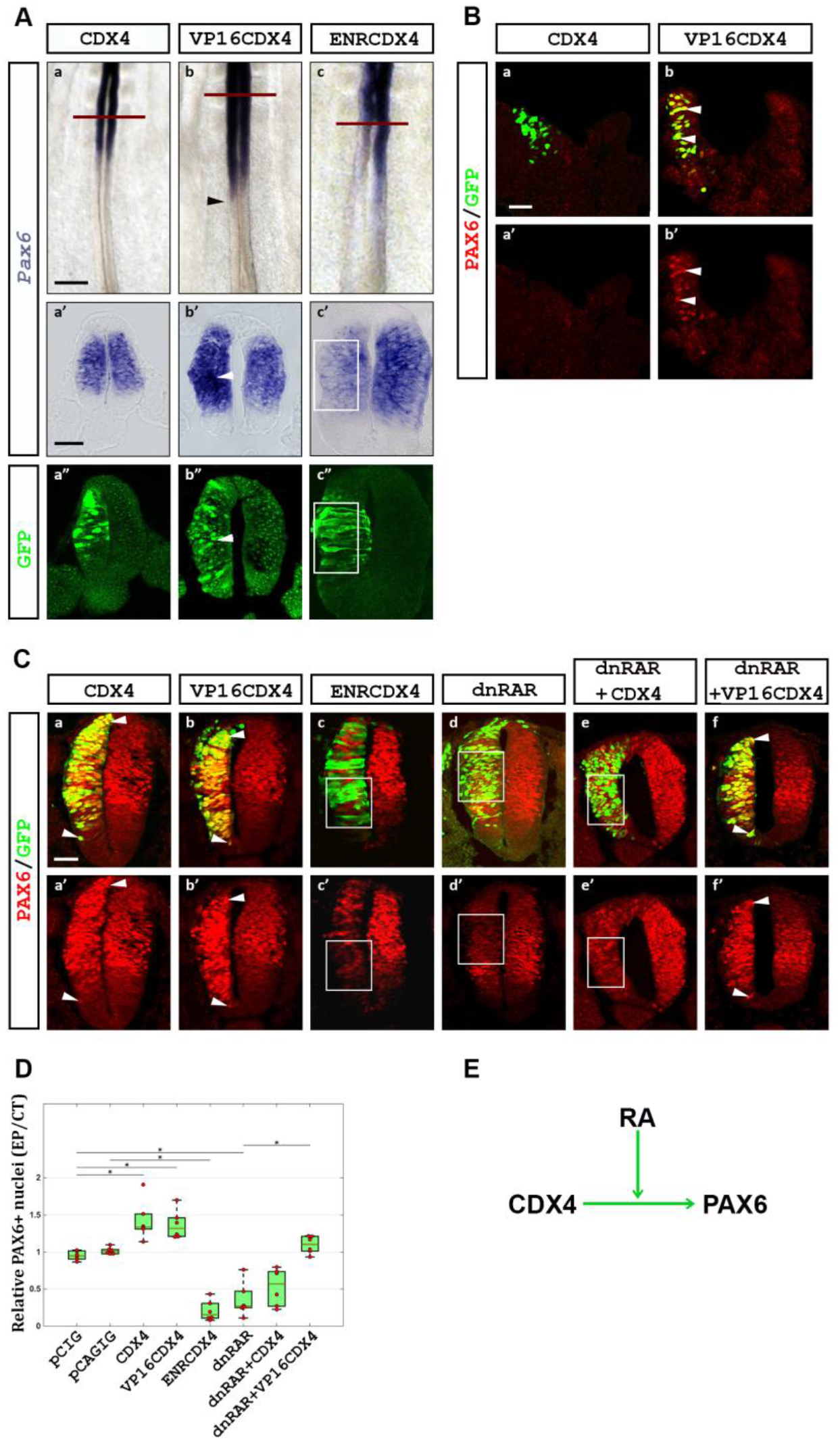
CDX4 activation of *Pax6* transcription is RA-dependent. (**A**) CDX4 regulates *Pax6* transcription in the rostral PNT. In the PNT, *Cdx4* has no effect (a, a’; n=6/6), *VP16Cdx4* induces (arrowheads in b, b’; n=4/6), and *EnRCdx4* downregulates (c, c’, box; n=6/6) *Pax6* transcription (purple signal by ISH; green GFP tag labeled by IHC). Embryos were electroporated at HH10-11 and analyzed 8 hpe (HH12-13). (**B**) Similarly, ectopic *VP16Cdx4* (n=3/4) but not *Cdx4* (n=0/4), causes ectopic PAX6 protein accumulation (arrowheads). (**C**) CDX4 requires Retinoic Acid (RA) to activate *Pax6* transcription in rostral regions. In the NT, both *Cdx4* and *VP16Cdx4* overexpression result in ectopic PAX6 protein accumulation (a, a’, b, b’; arrowheads; n=6/6 for both), whereas *EnRCdx4* overexpression causes the loss of PAX6 (c, c’; box; n=6/6). Inhibition of RA signaling using a dominant negative RA receptor (*dnRAR*) causes the loss of PAX6 (d, d’; box; n=6/6). In the absence of RA signaling, *Cdx4* overexpression is unable to induce ectopic PAX6 (e, e’; box; n=6/6). Under similar conditions, *VP16Cdx4* overexpression induces ectopic PAX6 (f, f; arrowheads; n=6/6). Embryos were electroporated at HH10-11 and analyzed 24 hpe (HH16-17). (**D**) Quantification of PAX6 positive cells after experiments shown in C. Box-scatter plot representing ratio of PAX6 positive cells on electroporated side to that on the contralateral control side (as per Karaz et al., 2016). Cells were counted using ImageJ. Significance is shown with a bar and a star (two tailed t-test analysis, p<0.05). (**E**) Summary of results. Scale bar is 200μm for whole mount and 40μm for transverse sections.

To test if CDX4 is necessary for *Pax6* activation in the PNT, we outcompeted endogenous CDX4 by overexpressing a dominant negative form of CDX4 in which the transcription activation domain of the protein was replaced with the transcriptional repressor domain of the *Drosophila* Engrailed protein (ENRCDX4; Han and Manley, 1993). This chimeric form of CDX4 has been shown to repress transcription of downstream CDX targets (e.g., *Hox* genes; Bel-Vialar et al., 2002; Isaacs et al., 1998). Overexpression of *EnRCdx4* caused *Pax6* downregulation in the rostral PNT (8 hpe; Fig 2Ac; n=6/6), indicating that in this region, CDX4 is necessary for *Pax6* transcription.

### CDX4 activation of *Pax6* in the PNT is dependent on Retinoic Acid signaling

Transcription of *Pax6* is restricted to the rostral PNT despite that *Cdx4* is transcribed in both caudal and rostral PNT regions (Fig 1D) and, upon overexpression, CDX4 can induce *Pax6* transcription ectopically (Fig 1F, 2A). To investigate the possible mechanisms that restrict *Pax6* transcription to the rostral PNT, we turned our attention to Retinoic Acid (RA). Somite-derived RA regulates spinal cord neurogenesis by activating numerous target genes in the rostral PNT, including *Pax6* (Novitch et al., 2003; Pituello et al., 1999). Given that RA and CDX4 interact during zebrafish spinal cord cell specification (Chang et al., 2016; Lee and Skromne, 2014), we hypothesized that RA and CDX4 might also interact during spinal cord maturation. To test this hypothesis, we electroporated PNT with dominant negative RA receptors (dnRAR) to block RA signaling (Novitch et al., 2003), and then analyzed the transcription of *Pax6* 24-hpe, at a time when electroporated cells would be undergoing maturation. As previously shown (Novitch et al., 2003), overexpression of *dnRAR* blocked RA signaling and caused *Pax6* down regulation (Fig 2Cd, D), even as *dnRAR* enhanced *Cdx4* transcription (Fig S2). To test if induction of *Pax6* by CDX4 is RA-dependent, we co-electroporated different *Cdx4* constructs together with *dnRAR*. In RA-deficient cells, CDX4 was unable to induce *Pax6* (Fig 2Ce, D; n=6/6), despite its ability to do so in RA-responsive cells (Fig 1D). Significantly, however, VP16CDX4 was able to induce *Pax6* transcription even in the absence of RA (Fig 2Cb, Cf, D; n=6/6). Together, these results suggest that *Pax6* activation by CDX4 is dependent on RA signaling, and illuminates a mechanism for the restricted transcription of *Pax6* to the rostral portion of the PNT (Fig 2E).

### CDX4 inhibits PAX6-dependent activation of *Ngn2* in the NT

PAX6 is present in both the rostral PNT and the NT, but it only activates neural differentiation genes in the NT (Bel-Vialar et al., 2007). Then, what prevents PAX6 from prematurely activating neural differentiation genes in the PNT? To address this question we analyzed the transcription of the neural differentiation gene *Ngn2,* a downstream target of PAX6 (Scardigli et al., 2003). *Ngn2* transcription domain is nested within that of *Pax6* and lays immediately rostral to that of *Cdx4* (Fig 1B; Fig 3A), raising the possibility that CDX4 activity is incompatible with *Ngn2* transcription. To test this possibility, we overexpressed *Cdx4, VP16Cdx4* and *EnRCdx4* in HH10-11 embryos and analyzed their effect on NGN2 distribution at HH16-17 (24 hpe). As expected, ENRCDX4 caused a loss of NGN2 (24 hpe; Fig 3Bc, 3C; n=6/6), as this construct also reduced the levels of PAX6 (24 hpe; Fig 2Cc), and PAX6 is required for *Ngn2* transcription (Scardigli et al., 2003). Surprisingly, CDX4 and VP16CDX4 also caused the loss of NGN2 (24 hpe; Fig 3Ba-b, 3C; n=6/6), under conditions that resulted in ectopic PAX6 (Fig 2Ca-b), suggesting that CDX4 represses *Ngn2.* This result was consistent with *Cdx4* and *Ngn2* complementary expression domains (Figs 1B, 3A). To confirm that CDX4 represses *Ngn2* in the presence of PAX6, we repeated the experiment by simultaneously co-expressing *Cdx4* and *Pax6.* While PAX6 on its own was able to ectopically activate *Ngn2* (Fig 3Bd, C; n=6/6; Bel-Vialar et al., 2007), it was unable do so in the presence of CDX4 (Fig 3Be, C; n=6/6). Taken together, these results suggest that CDX4 in the PNT promotes *Pax6* and prevents *Ngn2* transcription (Fig 3D).

**Fig 3.**
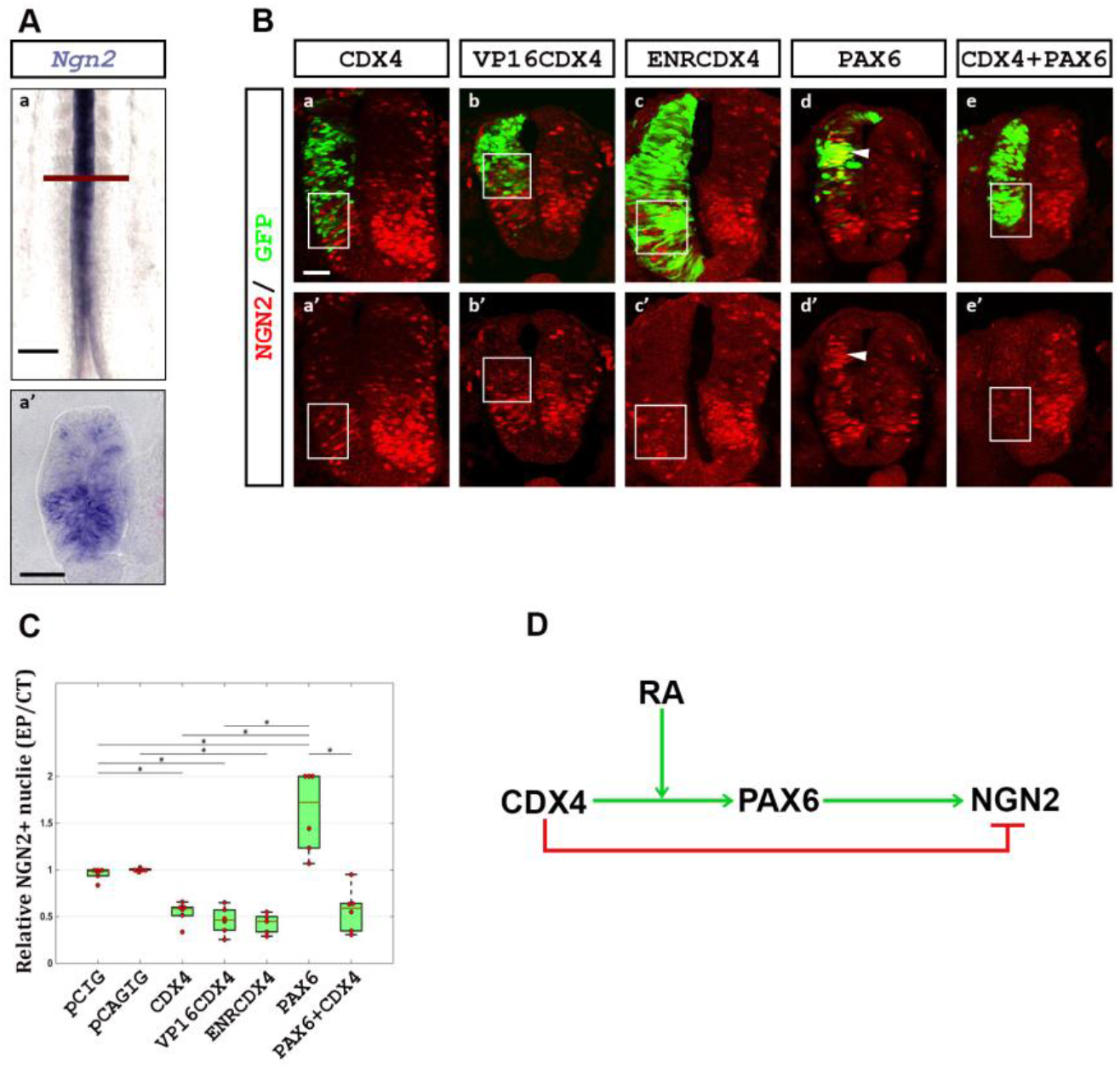
CDX4 inhibits early cell maturation by repressing the neural differentiation gene *Ngn2.* (**A**) *Ngn2* expression in the NT of wild type HH11 embryos. Expression is first observed caudally at the site of NT closure. (**B**) *Cdx4* and *VP16Cdx4* overexpression results in the loss of NGN2 (a, a’, b, b’; boxes; n=6/6 for both conditions), despite both inducing the Ngn2-activator *Pax6* (Fig 2A, C). *EnRCdx4* overexpression also causes the loss of NGN2 (c, c’; box; n=6/6), similar to its effect on PAX6 (Fig 2Cc). *Pax6* overexpression results in ectopic NGN2 (d, d’; arrowhead; n=6/6), but not in the presence of *Cdx4* (e, e’; box; n=6/6). Embryos were electroporated at HH10-11 and analyzed 24 hpe (HH16-17). (**C**) Quantification of NGN2 positive cells after experiments shown in B. Box-scatter plot representing ratio of NGN2 positive cells on electroporated side versus contralateral control side. Cells were counted using ImageJ. Significance is shown with a bar and a star (two tailed t-test analysis, p<0.05). (**D**) Figure summarizing CDX4-PAX6-NGN2 interactions. Scale bar is 200μm for whole mount and 40μm for transverse sections.

### CDX4 inhibits *Nkx1.2* expression in early neural progenitor cells

*Cdx4* transcription domain in the PNT includes the caudal region that contains *T/Bra-, Sox2+* and *Nkx1.2+* early neural progenitors. This observation prompted us to ask whether CDX4 also regulates aspects of early PNT maturation. To address this question, we electroporated the caudal PNT with different *Cdx4* constructs at HH10-11 and, after growing the embryos for 8 hpe to HH12-13, we analyzed the transcription of the early PNT marker *Nkx1.2* (Delfino-Machin et al., 2005; Gouti et al., 2017; Gouti et al., 2015; Sasai et al., 2014). Overexpression of *Cdx4* and *VP16Cdx4* caused downregulation of *Nkx1.2* transcription (Fig 4A; n=6/6 for both conditions), suggesting that CDX4 is a negative regulator of *Nkx1.2.* Unexpectedly, however, *EnRCdx4* overexpression also caused *Nkx1.2* downregulation (Fig 4A; n=6/6), suggesting that CDX4 is also necessary for *Nkx1.2* transcription. We interpret these results to indicate that *Nkx1.2* is under both positive and negative CDX4 regulation, with high levels of CDX4 repressing *Nkx1.2* (Fig 4C).

**Fig 4.**
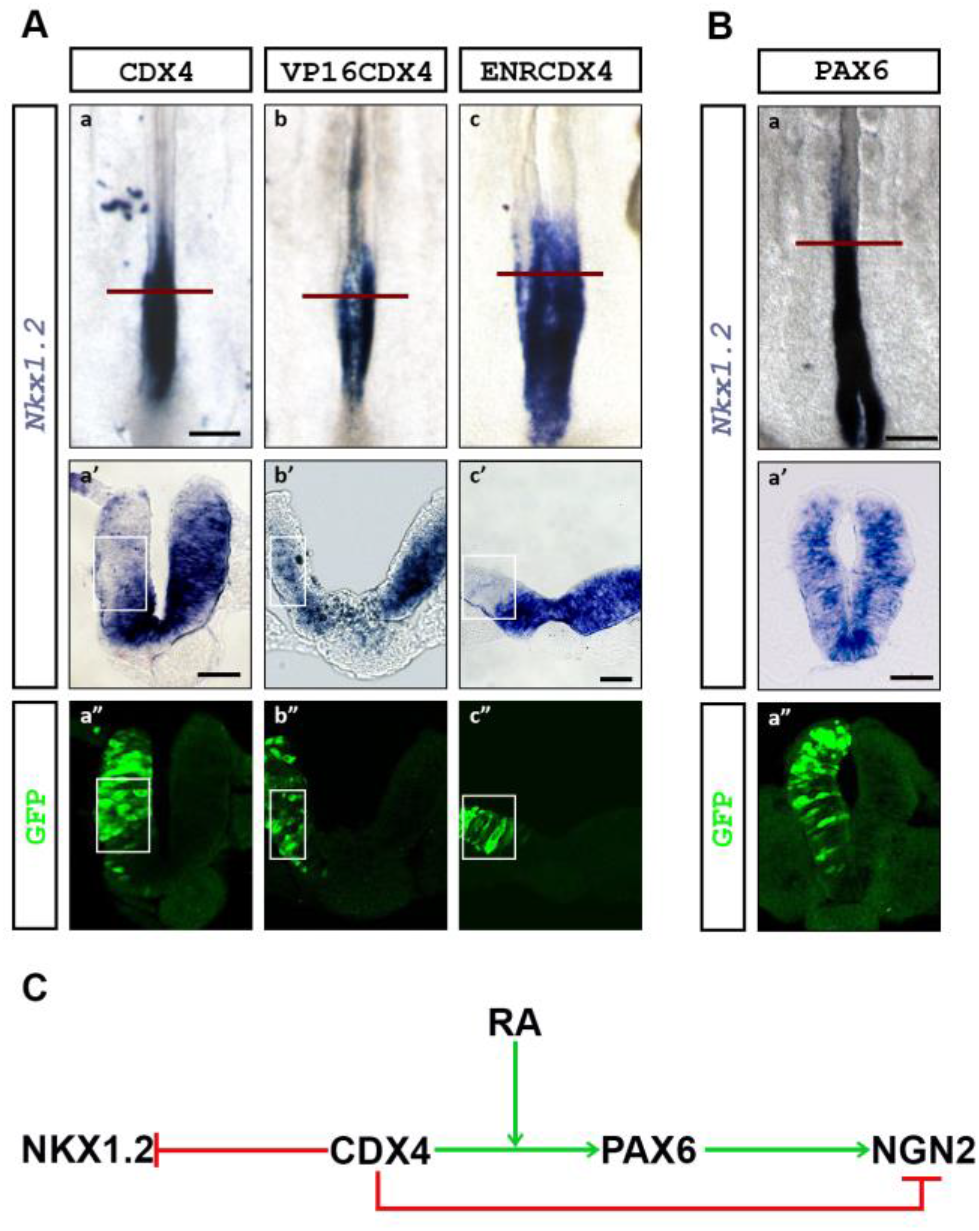
CDX4 regulates the transcription of Nkx1.2, an early neural progenitor cell identity marker. (**A**) Overexpression of *Cdx4* (a, a’), *VP16Cdx4* (b, b’), and *EnRCdx4* (c, c’) downregulate *Nkx1.2* transcription (boxed region; n=6/6 for all conditions). (**B**) Overexpression of *Pax6* has no effect on *Nkx1.2* transcription (n=6/6). Embryos were electroporated in the caudal PNT at HH10-11 and analyzed 8-hpe (HH12-13). (**C**) Figure summarizing CDX4-NKX1.2 interactions. Scale bar is 200μm for whole mount and 40μm for transverse sections.

One possible mechanism to explain the loss of *Nkx1.2* transcription after *Cdx4* overexpression is that CDX4 activates neural differentiation genes that then repress *Nkx1.2.* Such a candidate could be *Pax6,* as its expression domain and that of *Nkx1.2* are mutually exclusive (summarized in Fig 1A), its transcription is induced by CDX4 (Fig 2), and its overexpression induces neural differentiation (Bel-Vialar et al., 2007). To test this possibility, we repeated these experiments overexpressing *Pax6* and analyzing *Nkx1.2* transcript at HH12-13. Under these conditions, PAX6 did not change *Nkx1.2* transcription (Fig 4B; n=6/6). While this result does not rule out indirect mechanisms for the regulation of *Nkx1.2* by CDX4, the result suggests that PAX6 is not a *Nkx1.2* repressor (Fig 4C).

### NKX1.2 and PAX6 interactions result in the segregation of their expression domains to different regions of the PNT

*Nkx1.2* and *Pax6* transcription domains are mutually exclusive, but both span *Cdx4* transcription domain (summarized in Fig 1A), suggesting the possibility of cross-regulatory interactions between these genes. To test for possible cross-regulatory interactions between *Nkx1.2* and *Pax6,* and between these two genes and *Cdx4,* we analyzed the expression of these genes after their overexpression in the caudal PNT at HH10-11. To analyze NKX1.2 function, we overexpressed the mouse version of *Nkx1.2 (mNkx1.2),* which acts as a repressor in mouse cell lines (Tamashiro et al., 2012) and chicken embryos (Sasai et al., 2014). Consistent with previous report (Sasai et al., 2014), overexpression of *mNkx1.2* represses *Pax6* at HH12 (8 hpe; Fig 5Ac, n=6/6). In addition, mNKX1.2 also repressed *cNkx1.2* transcription (Fig 5Aa, n=6/6), suggesting negative autoregulation. However, *mNkx1.2* overexpression had no effect on *Cdx4* transcription (Fig 5Ab; n=6/6). Using the same strategy, we analyzed PAX6 activity on *Cdx4.* In this experiment, overexpression of *Pax6* downregulated and dominant-negative *EnRPax6* upregulated *Cdx4* transcription (Fig 5B; n=6/6 for all conditions), suggest that PAX6 functions to represses *Cdx4*. Together, these results providing a mechanism to explain the segregation of *Nkx1.2* and *Pax6* transcriptional domains to caudal and rostral PNT, respectively, and the downregulation of *Cdx4* in the caudal NT (Fig 5C; see model below).

**Fig 5.**
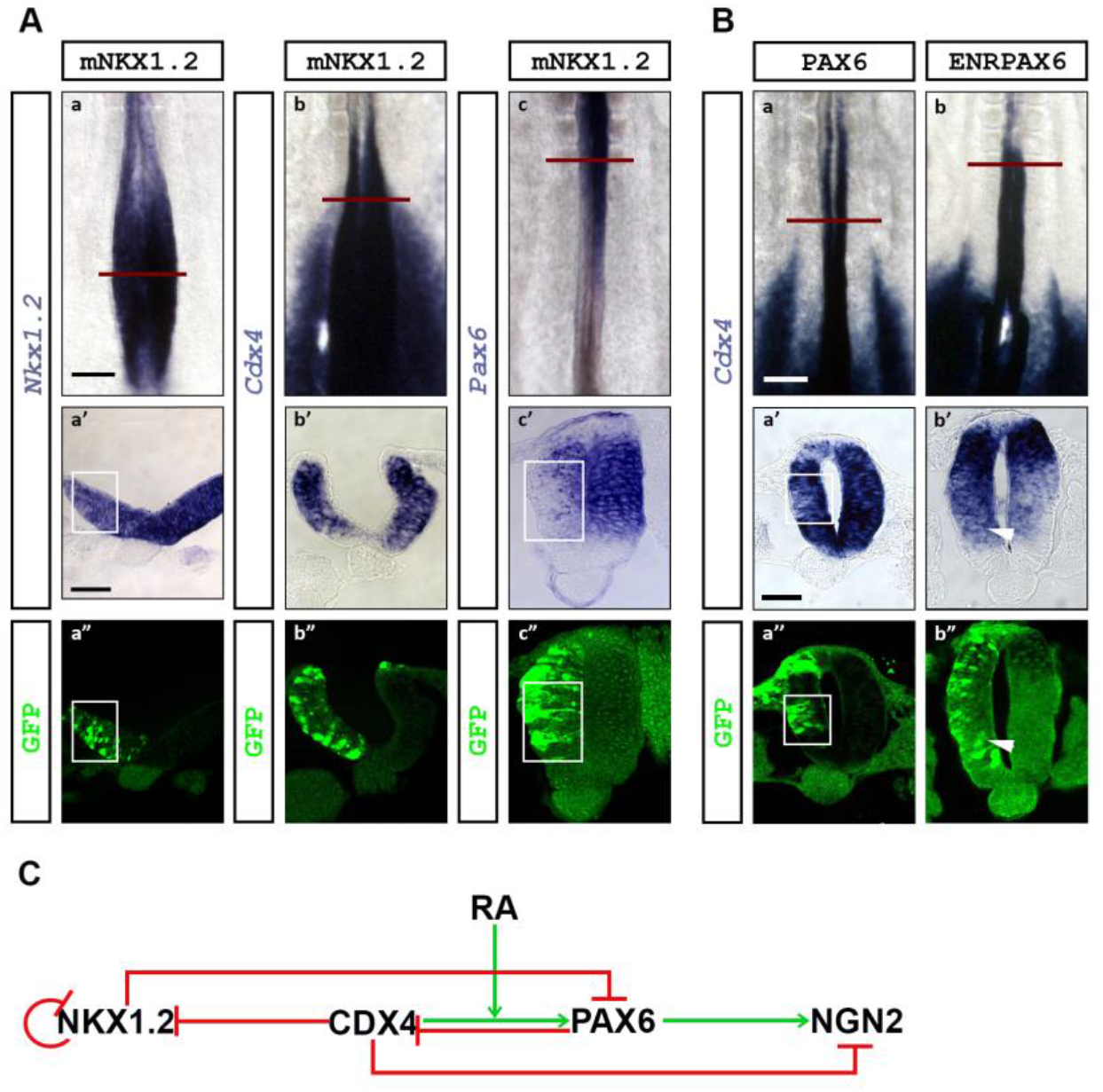
NKX1.2 and PAX6 contribution to *Cdx4* regulation. (**A**) NKX1.2 negatively regulates the transcription of its own gene and of *Pax6,* but not *Cdx4.* Overexpression of *mNkx1.2* downregulates *cNL·1.2* (a, a’) and *Pax6* (c, c’), without altering *Cdx4* transcription (a’, c’; boxed regions; n=6/6 for all conditions). (**B**) PAX6 represses *Cdx4.* Ectopic *Pax6* downregulates (a, a’; boxed region), and *EnRPax6* upregulates (b, b’; arrowhead) *Cdx4* transcription (n=6/6 for both conditions). Embryos were electroporated at HH10-11 and analyzed 8hpe (HH12-13). (**C**) Figure summarizing NKX1.2-CDX4-PAX6 interactions. Scale bar is 200μm for whole mount and 40μm for transverse sections.

## DISCUSSION

### A gene regulatory network controlling spinal cord neuronal maturation

#### CDX4 promotes loss of potency in the caudal pre-neural tube

Previous work has shown that CDX are key in the establishment and subsequent differentiation of NMPs into neural and mesodermal precursors by balancing the activity of Wnt3a, Fgf8 and RA signaling (Amin et al., 2016; Chawengsaksophak et al., 2004; Gouti et al., 2017). Mouse embryos deficient for all *Cdx* genes fail to develop post-occipital structures due to the premature differentiation of NMP cell (Amin et al., 2016; van Rooijen et al., 2012). The primary cause for this premature differentiation is the premature activation of the RA pathway, which in cell culture conditions causes NMPs to follow a neural fate by maintaining *Sox2* and *Nkx2.1,* and repressing *T/Bra* transcription (Gouti et al., 2017).

Here we show important additional functions for CDX4, the only CDX member present in chick PNT post HH12 (Marom et al., 1997), in the progressive maturation of spinal cord neuronal progenitors. As NMPs’ descendants acquire neural identity (from *T/Bra+, Sox2+, Nkx1.2+* to *T/Bra-, Sox2 +, Nkx1.2+),* CDX4 promote their further maturation by repressing *Nkx1.2* transcription (Fig 4). Control of *Nkx1.2* transcription is tightly balanced by Wnt and FGF signaling (Bertrand et al., 2000; Tamashiro et al., 2012), signals that are regulated by CDX (Chawengsaksophak et al., 2004; Gouti et al., 2017; Savory et al., 2009; van Rooijen et al., 2012). We speculate that increasing or decreasing CDX4 levels could cause an imbalance in Wnt and FGF that could lead to a loss of *Nkx1.2* transcription. Given that NKX1.2 inhibits floor plate cell specification by repressing *Pax6* (Sasai et al., 2014), we propose that CDX4 downregulation of *Nkx1.2* is one of the first steps in PNT cell maturation.

#### CDX4 promotes neural cell determination in the rostral pre-neural tube

Progression of cells from caudal to rostral PNT is marked by the acquisition of neural determination markers. CDX4 promotes new maturation states by directing *Pax6* activation, a factor involved in neural progenitor maturation (Bel-Vialar et al., 2007). CDX factors have been observed to increase *Pax6* transcription in embryoid bodies (McKinney-Freeman et al., 2008). We propose that in the PNT, CDX4 regulation of *Pax6* occurs via two distinct mechanism (Fig 6): by the indirect down regulation of the *Pax6* repressor NKX1.2 (Fig 5), and by the direct activation of *Pax6* transcription (Fig 2). Importantly, CDX4 activation of *Pax6* is dependent on RA (Fig 2), which is secreted from somites (Molotkova et al., 2005; Olivera-Martinez and Storey, 2007). This spatial distribution of RA restricts the *Pax6* inducing activity of CDX4 to the rostral PNT. RA/CDX regulation of *Pax6* is likely to be evolutionarily conserved across vertebrates, as in all species examined the second intron of *Pax6* contains an ultraconserved non-coding region that harbors both RA response elements (RAREs; Cunningham et al., 2016) and CDX4 binding sites (Paik et al., 2013 and Fig S3). In addition to DNA binding, RA has been implicated in opening up the *Pax6* locus by antagonizing FGF signaling (Patel et al., 2013). In this scenario, RA could function to provide locus accessibility to CDX4 and other factors. Thus, in the PNT, RA provides the context in which CDX4 can further promote neural progenitor cell maturation.

**Fig 6.**
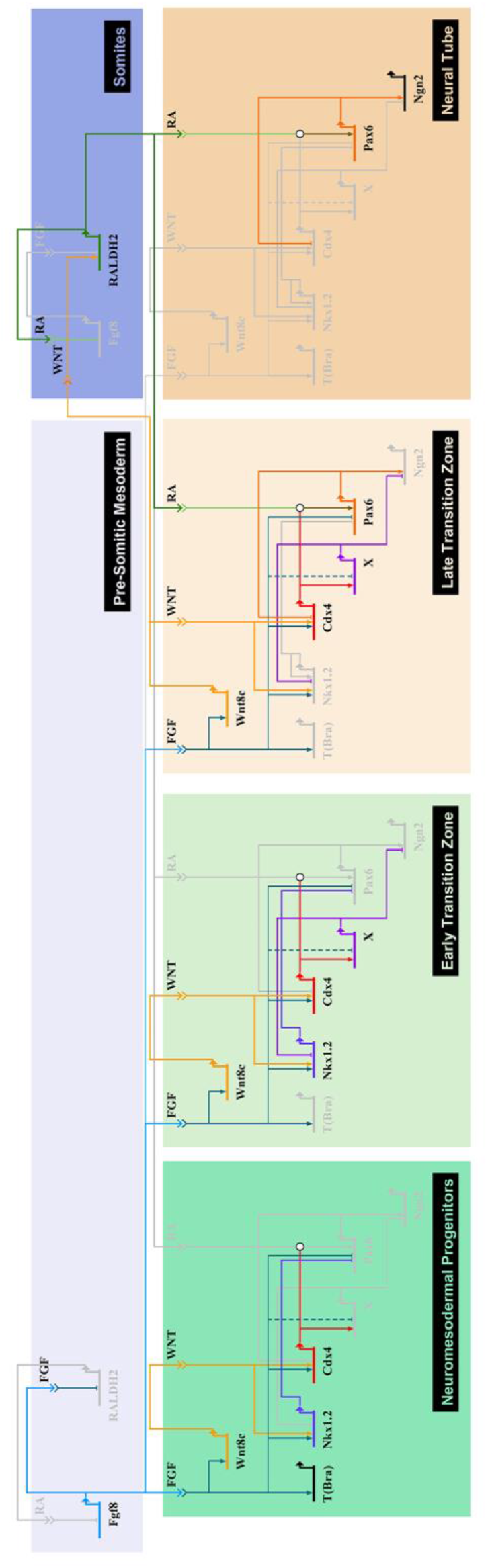
Proposed gene regulatory network controlling spinal cord neuronal cell maturation. Gene regulatory network of the genetic interactions identified in figures 1-5, superimposed to the FGF-Wnt8C-RA signaling network shown by others to regulate cell transitions states during spinal cord neuronal cell maturation (Olivera-Martinez and Storey, 2007). Network map was generated using Biotapestry (Longabaugh et al., 2005). In this model, CDX4 is at the core of the gene regulatory network that coordinates upstream signaling information into downstream transcriptional response.

#### CDX4 prevents premature neural cell differentiation

PAX6 induces neural cell differentiation (Bel-Vialar et al., 2007), and yet, despite *Pax6* being induced by CDX4 in the rostral PNT (Fig 2), neural cell differentiation does not begin until after the NT has formed and *Cdx4* has been down regulated (Fig 1). Two mechanisms by which PAX6 promotes differentiation is by downregulating *Cdx4* (Fig. 5B) and by activating *Ngn2* (Bel-Vialar et al., 2007; Scardigli et al., 2003), a gene that promotes cell cycle exit and further differentiation (Lacomme et al., 2012). Our data shows that CDX4 represses *Ngn2* transcription even in the presence of PAX6 (Fig 3), thus priming but delaying spinal cord terminal cell differentiation. Along the caudal-to-rostral axis of the neural tube, CDX4 transcription is gradually restricted to the dorsal neural tube (Fig 1), at a time when *Ngn2* transcription initiates ventrally (Fig 3). At the moment, it is unclear how CDX4 represses *Ngn2,* as CDX are known transcriptional activators (Isaacs et al., 1998). In sum, by regulating the activation of specification, determination and differentiation genes, CDX4 controls the transition of neural cells from one state to the next during the early maturation of the spinal cord. The regulation of cell transitions by CDX proteins may be a general property of this family of transcription factors, as CDX family members have also been described to control maturation of multipotent cell precursors in intestinal (Hryniuk et al., 2012; Saad et al., 2011) and hematopoietic (McKinney-Freeman et al., 2008; Wang et al., 2008) tissues.

### A model of spinal cord neuronal maturation that integrates transcription and signaling networks

In current models of spinal cord development, cells progressively lose potency and acquire neural characteristics under the control of FGF, Wnt and RA signaling (Diez del Corral and Storey, 2004; Gouti et al., 2014). Mutual interactions among these signaling factors restrict the activity of respective pathways to specific domains within the caudal and rostral PNT to direct cell fate decisions. In the caudal end, high levels of FGF promote *Wnt* transcription while repressing RA pathway activity through a variety of mechanisms (Boulet and Capecchi, 2012; Olivera-Martinez et al., 2012; Sakai et al., 2001; White et al., 2007). In turn, WNT8c promotes RA synthesis in anterior pre-somitic mesoderm and somites, away from the caudal domain of FGF activity. RA secreted from these anterior sources represses FGF synthesis, helping establish and refine the high-caudal to low-rostral gradients of FGF, and indirectly, Wnts (Diez del Corral et al., 2003; Kumar and Duester, 2014). This cross-repressive activities of FGF/Wnts and RA create a caudal-to-rostral gradient of potency signals and a rostral-to-caudal gradient of pro-differentiation signaling that promote the gradual maturation of spinal cord cells (Fig 1, 2). Molecularly, FGF and Wnt maintain NMPs cells by promoting the transcription of multipotency genes *T/Bra, Sox2* and *Nkx1.2,* while simultaneously repressing the differentiation genes *Pax6,* and *Ngn1/2.* In contrast, RA promotes differentiation by repressing *T/Bra* and *Nkx1.2* and inducing *Pax6* and *Ngn1/2* transcription. Thus, the switch from NMP to pre-neural to neurogenic identities is the response of cells to change in extracellular signals (Diez del Corral et al., 2003).

We have expanded the model of spinal cord neurogenesis by integrating signaling and transcription network models (Fig 6). The FGF-Wnt-RA network model provides a series of interactions that result in the spatiotemporal separation of regulatory inputs without providing intracellular mechanisms for the specification and separation of cells states, whereas the transcription factor network provides a molecular mechanism for the specification of different cellular states, but lacks the inputs necessary to drive the system forward. CDX4, at the core of the transcription factor network, provides an integration point for the inputs to regulate effector genes, as *Cdx4* transcription is directly regulated by FGF, Wnt and RA (Chang et al., 2016; Keenan et al., 2006; Lee and Skromne, 2014; Nordstrom et al., 2006; Tamashiro et al., 2012). FGF and Wnt promote potency by directly activating *Nkx1.2* (Diez del Corral et al., 2003; Tamashiro et al., 2012), but also initiate the loss of potency by sustaining *Cdx4* transcription that indirectly represses *Nkx1.2* (Fig 4B). A similar “dual-activity” phenomenon is observed in the regulation of *Pax6,* with FGF both repressing (Bertrand et al., 2000) and activating (via CDX4; Fig 2B) *Pax6* transcription. While the mechanism by which CDX4 antagonizes FGF activity at the *Pax6* locus is unknown, it may involve a change in *Pax6* chromatin state. FGF signaling has been shown to cause the translocation of the *Pax6* locus to the nuclear boundary associated with inactive chromatin (Patel et al., 2013). CDX4 could antagonize this activity, as CDX family members have been associated with the clearance of repressive histone modifications in other loci (e. g., *Hox;* Mazzoni et al., 2013). Regardless of the mechanism, we observe that for two genes, *Nkx1.2* and *Pax6,* CDX4 antagonizes FGF and synergizes with RA. We propose this FGF-Cdx/RA antagonism provide a time delay mechanism to separating early, intermediate and late states of cell differentiation. Experiments are under way to test the interactions between the signaling and transcription factors discussed in this model.

### CDX and the coordinated control of spinal cord neuronal maturation, patterning and growth

In addition to regulating spinal cord neuronal maturation, CDX factors are key regulators of axial patterning and elongation. In the context of patterning, CDX4 work together with FGF (and Wnts) to activate transcription of branchial and thoracic *Hox* genes (Bel-Vialar et al., 2002; Marletaz et al., 2015; Nordstrom et al., 2006; Shimizu et al., 2006; Skromne et al., 2007) and antagonizes RA’s ability to induce hindbrain *Hox* genes (Lee and Skromne, 2014; Marletaz et al., 2015; Skromne et al., 2007). Significantly this interaction is in contrast to the CDX4-FGF antagonism and CDX4-RA cooperation that we observed during spinal cord neuronal maturation (Fig 6). The molecular mechanism underlying this context-dependent switch in CDX4 activities is currently unknown. However, CDX4 involvement in both processes is significant as it provides a mechanism for coordinating the maturation and anterior-posterior identity specification of spinal cord neurons.

CDX role in vertebrate body extension involves maintaining progenitor population via two distinct mechanisms. Early in spinal cord development CDX cooperate with T/BRA to promote FGF and Wnt signaling cascades and sustain NMP proliferation (Amin et al., 2016; Gouti et al., 2017), whereas, later in development, CDX activate *Hox13* genes involved in axial termination (van de Ven et al., 2011; Young et al., 2009). Mutations in mouse that inactive *Cdx* or prematurely activate *Hox13* impairs elongation and morphogenesis of the spinal cord neuroepithelium, which results in irregular or duplicated neural structures (van de Ven et al., 2011). These neural tube defects are similar to those observed in mutants in the mesoderm specification genes *T/Bra* and *Tbx6* (Chapman and Papaioannou, 1998; Yamaguchi et al., 1999), leading to the proposal that caudal spinal cord defects associated with the loss of CDX arise through defects in the specification of NMP descendent (van de Ven et al., 2011). In light of our results, however, the neural tube abnormalities associated with CDX loss could also be explained, at least in part, to defects in spinal cord neuronal maturation. Future work will need to determine the contextual contribution of CDX in coordinating spinal cord cell maturation, differentiation and axial identity specification.

## MATERIALS AND METHODS

### Chicken embryo incubation and harvesting

Fertilized broiler chicken eggs (Morris Hatchery, Inc.; Miami, FL) were incubated at 38.2° C in a humid chamber until reaching the appropriate stage of development. The embryos were staged according to Hamburger and Hamilton normal table of development (Hamburger and Hamilton, 1951). Embryos post-electroporation were incubated until stipulated time for further analysis.

### DNA constructs and chicken *in ovo* electroporation

Gene overexpression studies were done using standard cloning and electroporation techniques. To achieve high level of gene expression and to track electroporated cells, gene of interest was cloned either into pCIG or pCAGIG vector (Matsuda and Cepko, 2004; Megason and McMahon, 2002). These vectors use the chicken *Actin* promoter to drive high gene expression levels, and carry a *GFP* gene as a reporter for transcription. Genes of interest were either cloned into vectors in the laboratory (*Cdx4, VP16Cdx4, EnRCdx4, mNkx1.2;* for details see supplementary material), or obtained already in the appropriate vector from other laboratories (Pax6-pCIG and EnRPax6-pCIG were kindly provided by Dr. Francois Medevielle (Bel-Vialar et al., 2007); and mNkx1.2-pEF2 was kindly provided by Dr. Yusuke Marikawa (Tamashiro et al., 2012). Plasmids for electroporation were purified using QIAGEN maxi-prep kit, and diluted to a final concentration of 0.5 μg/μl in 1X PBS, with 50ng/ml Fast Green dye to aid in the visualization of the cocktail mix during the procedure. Neural tube of chicken embryos stage HH10-11 were injected with the DNA cocktail mix and immediately electroporated unilaterally following standard protocols (Itasaki et al., 1999; Nakamura and Funahashi, 2001). Only normal-looking embryos with good electroporation in the desired region (e.g., neural tube, pre-neural tube, or caudal neural plate depending on experimental requirements) were selected for further processing by *in situ* hybridization or immunohistochemistry. Analysis was focused on same axial level in all stage: PNT for stage HH12-13 (prospective thoracic level; Liu et al., 2001), and NT for stage HH16-17 (thoracic level between somites 20-25; Evans, 2003).

### *In situ* hybridization

Analysis of gene transcription by *in situ* hybridization was done using digoxigenin (DIG)-labeled antisense RNA probes synthesized and hybridized using standard protocol (Wilkinson and Nieto, 1993). Briefly, embryos were harvested at the appropriate stage and fixed with 4% paraformaldehyde diluted in 1x PBS at 4° C overnight, before processing for *in situ* hybridization. After a series of washes, embryos were exposed overnight in hybridization solution to DIG-labeled antisense RNA probes against *Pax6, Nkx1.2, T/Bra,* or *Cdx4.* mRNA expression was detected using an Alkaline Phosphatase coupled Anti-DIG antibody (Roche) and developing embryos with nitro-blue tetrazolium salt (NBT, Thermo Scientific) and 5-bromo-4-chloro-3-indolyl-phosphate (BCIP, Biosynth) at room temperature until dark purple precipitate deposited revealing the areas of gene transcription. For double in situ hybridization, one of the probes was labeled with FITC and developed with Fast Red (Sigma-Aldrich). Post-development, embryos were washed with 1x TBST and then fixed in 4% PFA.

### Cryo-sectioning and Immunohistochemistry

Embryos harvested for immunohistochemistry (IHC) analysis were fixed with 4 *%* PFA for 3 hours at room temperature. Embryos were then embedded in Shandon M1 embedding matrix media (Thermo Scientific) and quickly frozen over dry ice. Mounted embryos were sectioned on Leica CM1850 cryostat and consecutive 20 μm thick sections were collected on positive-charged glass slides (Globe scientific). Antibody staining was performed following standard protocols on slides stacked in Shandon Sequenza slide rack (Thermo Scientific) and supported by Shandon cover plates.

Primary antibodies against anti-mouse PAX6, PAX7 and NKX6.1 were obtained from development Studies Hybridoma Bank. Anti-chicken NGN2 antibody was a kind gift from Dr. Bennett Novitch (Skaggs et al., 2011). Rabbit polyclonal antibody against GFP Tag was obtained from AnaSpec Inc. Goat anti-mouse Alexa Flour 488, Alexa Flour 556 and goat anti-guinea pig Alexa Flour 568 secondary antibodies (Invitrogen) were used for detecting primary antibodies. Sections were covered with DAPI-containing mounting media (Vectashield) and a cover slip, and sealed with nail polish.

### Microscopy

Whole embryo images were taken on Zeiss V20 Stereo microscope with an AxioCam MRc digital color camera (Carl Zeiss). Images of transverse section of neural tube were taken on AXIO Examiner Z1 compound microscope with an AxioCam MRc color camera (Carl Zeiss), or on a Leica SP5 confocal microscope (Leica). Confocal images, thickness 2.304 μm, were processed with ImageJ (Schneider et al., 2012). Images were processed for figures using Adobe Photoshop (CC2017, Adobe) for size and resolution adjustment, and for figure preparation.

### Quantification of IHC data

To quantify changes in the levels of candidate proteins after electroporation, cells positive for PAX6 or NGN2 were counted on both electroporated and control sides at the same dorsal-ventral position, and their relative ratio was determined. Images were processed with ImageJ IHC toolbox plugin (Shu et al., 2013) before cell counting to select for cells above threshold level as determined by the program algorithm. A total of 6 embryos per conditions were used for determining significance. Significance of difference between mean values of compared pairs was evaluated using two-tailed t-test (Microsoft Excel). Data for each condition was graphed into a box-plus-scatter plot using MATLAB (2014b, The MathWorks Inc., Natick, MA, 2014).

## AUTHOR CONTRIBUTIONS

P.J. and I.S. designed the experiments. P.J. performed the experiments. A. J. D. provided intellectual contributions towards designing and troubleshooting experiments. P.J. and I.S. analyzed the results. P.J., A.J.D and I.S. wrote the manuscript.

## ACKNOWLEDGEMENTS

We thank Dr. Ann Foley, Dr. Pantelis Tsoulfas and two anonymous reviewers for their expert insights in improving the quality of the original manuscript, and members of the Skromne lab for intellectual discussion, particularly Dr. S. Bandopadhyay. We also thank Dr. K. G. Story (U Dundee, UK), Dr. M. Gouldin (Salk Institute, USA). Dr. F. Medeville (CBI, France), Dr. S. Mackem (NCI, USA), Dr. Y. Marikawa (U Hawaii, USA), Dr. A. V. Morales (Cajal Institute, Spain) and Dr. B. Novitch (UCLA, USA) for generously providing essential constructs and antibodies.

## Supplemental information

### Supplemental materials and methods

Gene and gene constructs employed in this work where either obtained from other laboratories, or generated by us using standard molecular biology techniques and publicly available annotated sequences. A list of genes and constructs obtained from other laboratories is summarized in Table S1. A list of primers for genes and constructs generated by us is summarized in Table S2.

*Full length Cdx4 for in situ hybridization and sub-cloning (Cdx4-pGEM-T-Easy).* Full length *Cdx4* (NM_204614.1) was cloned from reverse transcribed, total mRNA from chicken embryos at different stages of development (HH4-HH12; qScript cDNA Synthesis kit, Quantabio), using primers designed with the online program Primer BLAST (Table S2). Fragment product of the correct size was then cloned using pGem-T Easy Plasmid (Promega,). Cloning of the correct gene was confirmed by sequencing. This construct was used to generate *in situ* hybridization probe and as a template for additional construct.

*Full length Cdx4 for chicken electroporation (Cdx4-pCIG).* Full-length chicken *Cdx4* was digested with SpeI and blunt ended with Mung Bean nuclease (NEB). Purified, linear *Cdx4* was then digested with EcoRI. The purified *Cdx4* fragment was then subcloned into pCIG previously digested with EcoRI-SmaI. This construct was used for overexpressing wild type *Cdx4* in chicken embryos by electroporation. pCIG contains nuclear GFP under IRES promoter for concomitant expression of GFP in electroporated cells (Megason and McMahon, 2002).

*Constitutively active Cdx4 for chicken electroporation (VP16Cdx4-pCIG).* The transactivator domain of the *VP16* was amplified from VP16-pCS2+ and fused to the C-terminal domain of Cdx4 containing the DNA binding homeodomain (corresponding to amino acids 119-364; renamed Cdx4-HD). Primers used for these amplifications are described in Table S2. Chimeric *VP16Cdx4* was then generated by PCR amplification from a mixture containing *VP16* and CdX4-HD fragments and VP16 forward and Cdx4-HD reverse primer. The segment was cloned into pGEM-T-easy and open reading frame confirmed by sequencing. *VP16Cdx4* was then digested using ClaI-EcoRI and inserted into ClaI-EcoRI sites of pCIG.

*Dominant negative Cdx4 for chicken electroporation (EnRCdx4-pCAGIG).* Engrailed (*EnR*) repressor domain from EnR-pCS2+ was digested with XhoI and blunt ended with Mung Bean nuclease. After purification, the fragment was digested with EcoRI and re-purified. This EcoRI-blunt *EnR* product was ligated to a blunt ended Cdx4 fragment generated using SmaI (nucleotide site 328). As a final step, the chimeric construct was ligated to pCAGIG vector digested with EcoRI-EcoRV. Several clones were analyzed by sequencing to confirm correct orientation of the *EnR* and *Cdx4* fragments, and the continuity of the open reading frame. pCAGIG contains GFP under IRES promoter for concomitant expression of cytoplasmic GFP in electroporated cells. pCIG is derived from pCAGIG backbone, with addition of nuclear localization signal in from of GFP, making to GFP concentrate in nucleus (Matsuda and Cepko, 2004).

*Full length mNkx1.2 for chicken electroporation (mNkx1.2 pCIG).* Mouse *Nkx1.2* was PCR amplified from the mNkx1.2-myctag pEf2 construct (gift from Y. Marikawa), using the primers shown in Table S2. The cloned segment was digested with ClaI and EcoRI included in the forward and reverse primers, respectively. Purified segment was then cloned into ClaI-EcoRI site of pCIG.

**Table S1.**
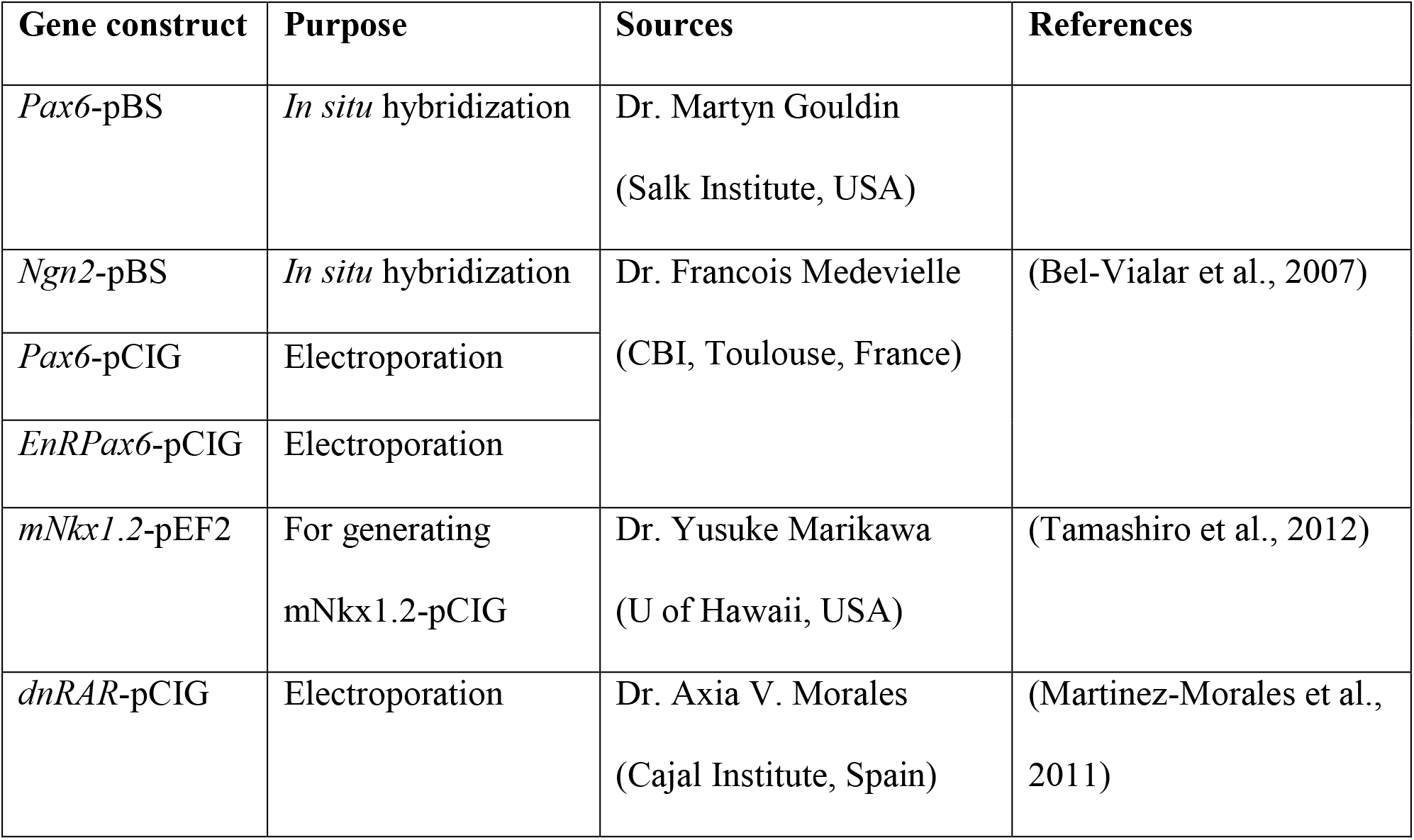
Genes and constructs received from other labs.

**Table S2.**
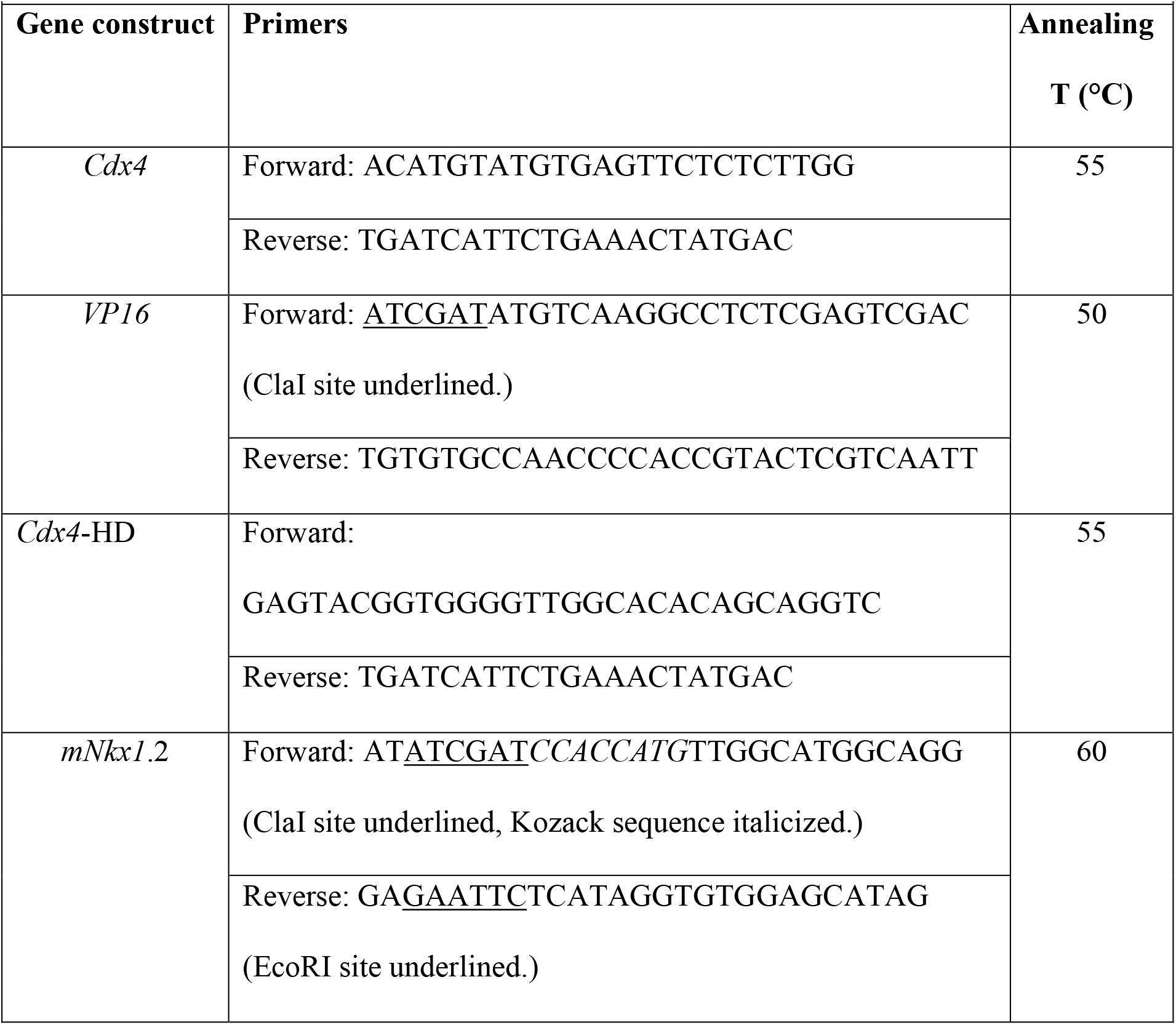
Genes cloned in the lab with respective primers.

### Supplementary figures

**Fig S1.**
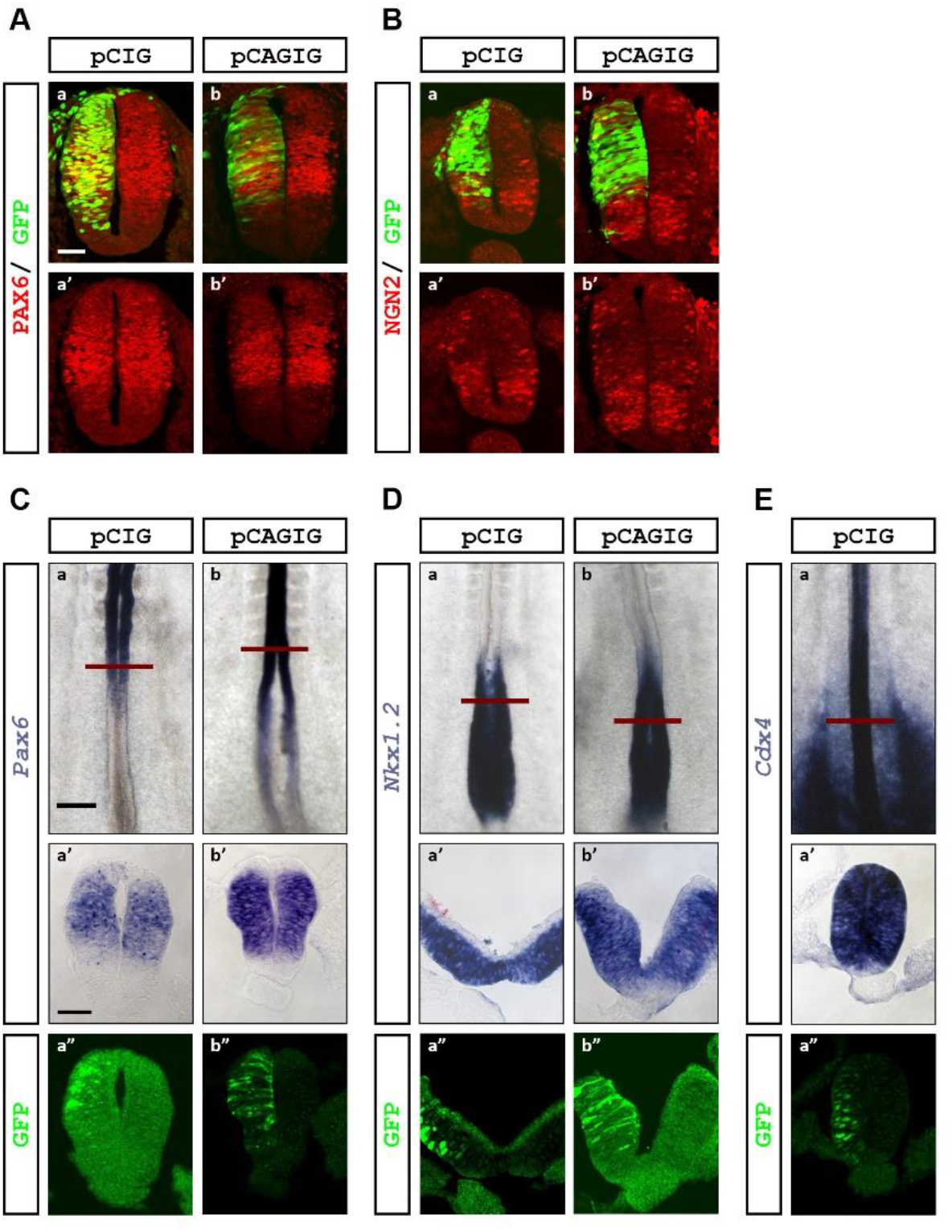
GFP overexpression does not affect wild type gene transcription of selected factors. pCIG (nuclear GFP) and pCAGIG (cytoplasmic GFP) ectopic expression did not change *Pax6* (A: IHC analysis, C: ISH analysis), NGN2 (B), *Nkx1.2* (D) and *Cdx4* (E) expression compared to contralateral control side (n=6/6).*Cdx4* expression analysis for pCAGIG over-expression was not done as none of the pCAGIG backbone construct were analyzed for *Cdx4* expression. Embryos were electroporated at H10-11 and analyzed at HH12-13 (8 hpe; ISH) or HH16-17 (24 hpe; IHC). Red bar shows RC level of transverse section. Scale bar is 200μm for whole mount and 40μm for transverse sections.

**Fig S2.**
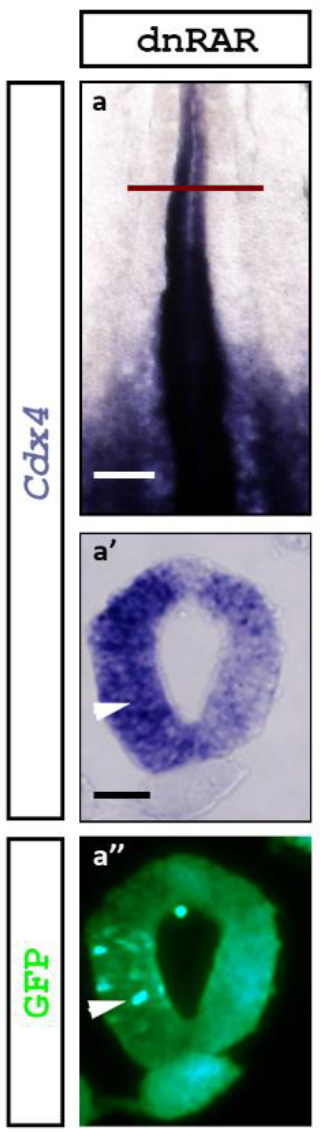
Loss of RA signaling causes expansion of *Cdx4* transcription domain in the NT. Expression analysis of *Cdx4* in embryos over expressing dnRAR. Arrowhead indicates electroporated side. Control non-electroporated side shows *Cdx4* downregulation (as seen in Fig. 1B). Embryos were electroporated at HH10-11 and analyzed at HH12-13 (8 hpe). Scale bar is 200μm for whole mount and 40μm for transverse section.

**Fig S3.**
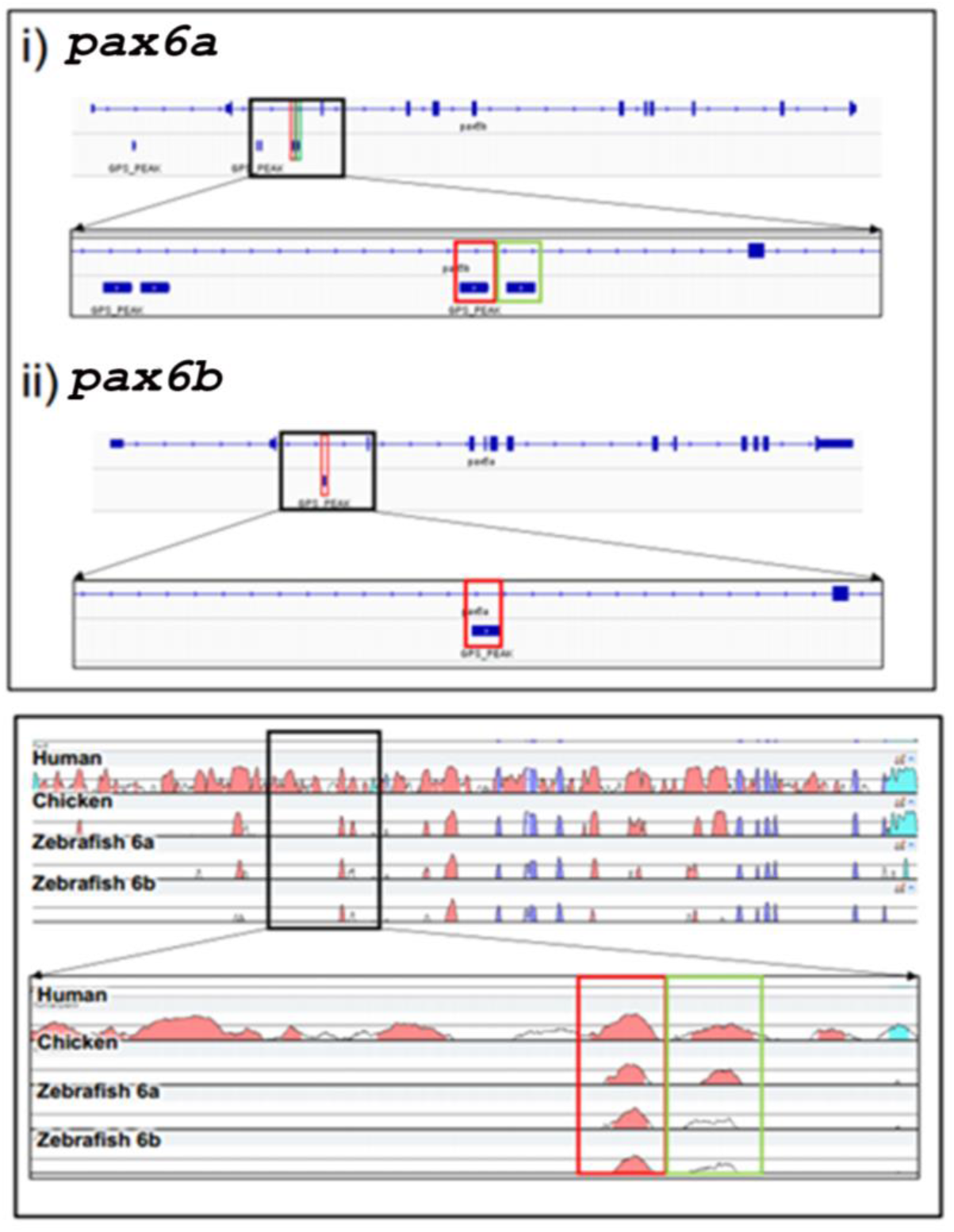
*Pax6* gene harbors evolutionarily conserved CDX4 binding sites. (Top) CDX4 ChIP-seq data showing CDX4 binding regions in zebrafish *pax6a* and *pax6b* genes (highlighted with green and red boxes; data obtained from Paik et al., 2013). (Bottom) Non-coding regions of zebrafish*pax6a* containing Cdx4 binding sites are conserved in human and chicken *Pax6* (highlighted green and red boxes are the same regions highlighted in the top panel).

## REFERENCES

1. Akai, J., Halley, P.A., Storey, K.G., 2005. FGF-dependent Notch signaling maintains the spinal cord stem zone. Genes Dev. 19, 2877–2887.

2. Amin, S., Neijts, R., Simmini, S., van Rooijen, C., Tan, S.C., Kester, L., van Oudenaarden, A., Creyghton, M.P., Deschamps, J., 2016. Cdx and T Brachyury Co-activate Growth Signaling in the Embryonic Axial Progenitor Niche. Cell Rep. 17, 3165–3177.

3. Bel-Vialar, S., Itasaki, N., Krumlauf, R., 2002. Initiating Hox gene expression: in the early chick neural tube differential sensitivity to FGF and RA signaling subdivides the HoxB genes in two distinct groups. Development. 129, 5103–5115.

4. Bel-Vialar, S., Medevielle, F., Pituello, F., 2007. The on/off of Pax6 controls the tempo of neuronal differentiation in the developing spinal cord. Dev. Bio. 305, 659–673.

5. Bertrand, N., Medevielle, F., Pituello, F., 2000. FGF signalling controls the timing of Pax6 activation in the neural tube. Development. 127, 4837–4843.

6. Boulet, A.M., Capecchi, M.R., 2012. Signaling by FGF4 and FGF8 is required for axial elongation of the mouse embryo. Dev. Bio. 371, 235–245.

7. Briscoe, J., Pierani, A., Jessell, T.M., Ericson, J., 2000. A homeodomain protein code specifies progenitor cell identity and neuronal fate in the ventral neural tube. Cell 101, 435–445.

8. Brown, J.M., Storey, K.G., 2000. A region of the vertebrate neural plate in which neighbouring cells can adopt neural or epidermal fates. Curr. Bio.10, 869–872.

9. Butler, S.J., Bronner, M.E., 2015. From classical to current: analyzing peripheral nervous system and spinal cord lineage and fate. Dev. Bio. 398, 135–146.

10. Cambray, N., Wilson, V., 2007. Two distinct sources for a population of maturing axial progenitors. Development. 134, 2829–2840.

11. Chang, J., Skromne, I., Ho, R.K., 2016. CDX4 and retinoic acid interact to position the hindbrain-spinal cord transition. Dev. Bio. 410, 178–189.

12. Chapman, D.L., Papaioannou, V.E., 1998. Three neural tubes in mouse embryos with mutations in the T-box gene Tbx6. Nature. 391, 695–697.

13. Chawengsaksophak, K., de Graaff, W., Rossant, J., Deschamps, J., Beck, F., 2004. Cdx2 is essential for axial elongation in mouse development. PNAS. 101, 7641–7645.

14. Cunningham, T.J., Colas, A., Duester, G., 2016. Early molecular events during retinoic acid induced differentiation of neuromesodermal progenitors. Bio. Open. 5, 1821–1833.

15. Davidson, E.H., 2006. The Regulatory Genome. Academic Press, Burlington.

16. Davidson, E.H., Levine, M.S., 2008. Properties of developmental gene regulatory networks. PNAS. 105, 20063–20066.

17. Delfino-Machin, M., Lunn, J.S., Breitkreuz, D.N., Akai, J., Storey, K.G., 2005. Specification and maintenance of the spinal cord stem zone. Development. 132, 4273–4283.

18. Deschamps, J., van den Akker, E., Forlani, S., De Graaff, W., Oosterveen, T., Roelen, B., Roelfsema, J., 1999. Initiation, establishment and maintenance of Hox gene expression patterns in the mouse. The Int. J. of Dev. Bio. 43, 635–650.

19. Deschamps, J., van Nes, J., 2005. Developmental regulation of the Hox genes during axial morphogenesis in the mouse. Development. 132, 2931–2942.

20. Diez del Corral, R., Olivera-Martinez, I., Goriely, A., Gale, E., Maden, M., Storey, K., 2003. Opposing FGF and retinoid pathways control ventral neural pattern, neuronal differentiation, and segmentation during body axis extension. Neuron. 40, 65–79.

21. Diez del Corral, R., Storey, K.G., 2004. Opposing FGF and retinoid pathways: a signalling switch that controls differentiation and patterning onset in the extending vertebrate body axis. BioEssays. 26, 857–869.

22. Evans, D.J., 2003. Contribution of somitic cells to the avian ribs. Dev. Bio. 256, 114–126.

23. Faas, L., Isaacs, H.V., 2009. Overlapping functions of Cdx1, Cdx2, and Cdx4 in the development of the amphibian Xenopus tropicalis. Dev. Dyn. 238, 835–852.

24. Gaunt, S.J., Drage, D., Trubshaw, R.C., 2005. cdx4/lacZ and cdx2/lacZ protein gradients formed by decay during gastrulation in the mouse. Int. J. Dev. Bio. 49, 901–908.

25. Gouti, M., Delile, J., Stamataki, D., Wymeersch, F.J., Huang, Y., Kleinjung, J., Wilson, V., Briscoe, J., 2017. A gene regulatory network balances neural and mesoderm specification during vertebrate trunk development. Dev. Cell. 41, 243–261.e247.

26. Gouti, M., Metzis, V., Briscoe, J., 2015. The route to spinal cord cell types: a tale of signals and switches. Trends Gen. 31, 282–289.

27. Gouti, M., Tsakiridis, A., Wymeersch, F.J., Huang, Y., Kleinjung, J., Wilson, V., Briscoe, J., 2014. In vitro generation of neuromesodermal progenitors reveals distinct roles for wnt signalling in the specification of spinal cord and paraxial mesoderm identity. PLoS Bio. 12, e1001937.

28. Hamburger, V., Hamilton, H.L., 1951. A series of normal stages in the *Development* of the chick embryo. J. Morph. 88, 49–92.

29. Han, K., Manley, J.L., 1993. Functional domains of the Drosophila Engrailed protein. EMBO J. 12, 2723–2733.

30. Hayward, A.G. 2nd, Joshi, P., Skromne, I., 2015. Spatiotemporal analysis of zebrafish hox gene regulation by Cdx4. Dev. Dyn. 244, 1564–1573.

31. Henrique, D., Abranches, E., Verrier, L., Storey, K.G., 2015. Neuromesodermal progenitors and the making of the spinal cord. Development. 142, 2864–2875.

32. Hryniuk, A., Grainger, S., Savory, J.G., Lohnes, D., 2012. Cdx function is required for maintenance of intestinal identity in the adult. Dev. Bio. 363, 426–437.

33. Isaacs, H.V., Pownall, M.E., Slack, J.M., 1998. Regulation of Hox gene expression and posterior development by the Xenopus caudal homologue Xcad3. EMBO J. 17, 3413–3427.

34. Itasaki, N., Bel-Vialar, S., Krumlauf, R., 1999. ‘Shocking’ developments in chick embryology: electroporation and in ovo gene expression. Nat. Cell Bio. 1, E203–207.

35. Karaz, S., Courgeon, M., Lepetit, H., Bruno, E., Pannone, R., Tarallo, A., Thouze, F., Kerner, P., Vervoort, M., Causeret, F., Pierani, A., D’Onofrio, G., 2016. Neuronal fate specification by the Dbx1 transcription factor is linked to the evolutionary acquisition of a novel functional domain. EvoDevo. 7, 18.

36. Keenan, I.D., Sharrard, R.M., Isaacs, H.V., 2006. FGF signal transduction and the regulation of Cdx gene expression. Dev. Bio. 299, 478–488.

37. Kumar, S., Duester, G., 2014. Retinoic acid controls body axis extension by directly repressing Fgf8 transcription. Development. 141, 2972–2977.

38. Lacomme, M., Liaubet, L., Pituello, F., Bel-Vialar, S., 2012. NEUROG2 drives cell cycle exit of neuronal precursors by specifically repressing a subset of cyclins acting at the G1 and S phases of the cell cycle. Mol. Cell. Bio. 32, 2596–2607.

39. Lee, K., Skromne, I., 2014. Retinoic acid regulates size, pattern and alignment of tissues at the head-trunk transition. Development. 141, 4375–4384.

40. Levine, M., Davidson, E.H., 2005. Gene regulatory networks for development. PNAS. 102, 4936–4942.

41. Liu, J.P., Laufer, E., Jessell, T.M., 2001. Assigning the positional identity of spinal motor neurons: rostrocaudal patterning of Hox-c expression by FGFs, Gdf11, and retinoids. Neuron. 32, 997–1012.

42. Lohnes, D., 2003. The Cdx1 homeodomain protein: an integrator of posterior signaling in the mouse. BioEssays. 25, 971–980.

43. Longabaugh, W.J., Davidson, E.H., Bolouri, H., 2005. Computational representation of developmental genetic regulatory networks. Dev. Bio. 283, 1–16.

44. Marletaz, F., Maeso, I., Faas, L., Isaacs, H.V., Holland, P.W., 2015. Cdx ParaHox genes acquired distinct developmental roles after gene duplication in vertebrate evolution. BMC Bio. 13, 56.

45. Marom, K., Shapira, E., Fainsod, A., 1997. The chicken caudal genes establish an anterior-posterior gradient by partially overlapping temporal and spatial patterns of expression. Mech. of Dev. 64, 41–52.

46. Matsuda, T., Cepko, C.L., 2004. Electroporation and RNA interference in the rodent retina in vivo and in vitro. PNAS.101, 16–22.

47. Mazzoni, E.O., Mahony, S., Peljto, M., Patel, T., Thornton, S.R., McCuine, S., Reeder, C., Boyer, L.A., Young, R.A., Gifford, D.K., Wichterle, H., 2013. Saltatory remodeling of Hox chromatin in response to rostrocaudal patterning signals. Nat. Neurosci. 16, 1191–1198.

48. McKinney-Freeman, S.L., Lengerke, C., Jang, I.H., Schmitt, S., Wang, Y., Philitas, M., Shea, J., Daley, G.Q., 2008. Modulation of murine embryonic stem cell-derived CD41+c-kit+ hematopoietic progenitors by ectopic expression of Cdx genes. Blood. 111, 4944–4953.

49. Megason, S.G., McMahon, A.P., 2002. A mitogen gradient of dorsal midline Wnts organizes growth in the CNS. Development. 129, 2087–2098.

50. Molotkova, N., Molotkov, A., Sirbu, I.O., Duester, G., 2005. Requirement of mesodermal retinoic acid generated by Raldh2 for posterior neural transformation. Mech. Dev. 122, 145–155.

51. Morales, A.V., de la Rosa, E.J., de Pablo, F., 1996. Expression of the cCdx-B homeobox gene in chick embryo suggests its participation in rostrocaudal axial patterning. Dev. Dyn. 206, 343–353.

52. Nakamura, H., Funahashi, J., 2001. Introduction of DNA into chick embryos by in ovo electroporation. Methods. 24, 43–48.

53. Nordstrom, U., Maier, E., Jessell, T.M., Edlund, T., 2006. An early role for WNT signaling in specifying neural patterns of Cdx and Hox gene expression and motor neuron subtype identity. PLoS Bio. 4, e252.

54. Novitch, B.G., Chen, A.I., Jessell, T.M., 2001. Coordinate regulation of motor neuron subtype identity and pan-neuronal properties by the bHLH repressor Olig2. Neuron. 31, 773–789.

55. Novitch, B.G., Wichterle, H., Jessell, T.M., Sockanathan, S., 2003. A requirement for retinoic acid-mediated transcriptional activation in ventral neural patterning and motor neuron specification. Neuron. 40, 81–95.

56. Olivera-Martinez, I., Harada, H., Halley, P.A., Storey, K.G., 2012. Loss of FGF-dependent mesoderm identity and rise of endogenous retinoid signalling determine cessation of body axis elongation. PLoS Bio. 10, e1001415.

57. Olivera-Martinez, I., Storey, K.G., 2007. Wnt signals provide a timing mechanism for the FGF-retinoid differentiation switch during vertebrate body axis extension. Development. 134, 2125–2135.

58. Paik, E.J., Mahony, S., White, R.M., Price, E.N., Dibiase, A., Dorjsuren, B., Mosimann, C., Davidson, A.J., Gifford, D., Zon, L.I., 2013. A cdx4-sall4 regulatory module controls the transition from mesoderm formation to embryonic hematopoiesis. Stem Cell. Rep. 1, 425–436.

59. Patel, N.S., Rhinn, M., Semprich, C.I., Halley, P.A., Dolle, P., Bickmore, W.A., Storey, K.G., 2013. FGF signalling regulates chromatin organisation during neural differentiation via mechanisms that can be uncoupled from transcription. PLoS Gen. 9, e1003614.

60. Peter, I.S., Davidson, E.H., 2013. Chapter 11 - Transcriptional Network Logic: The Systems Biology of Development A2 - Dekker, A.J. Marian WalhoutMarc VidalJob, Handbook of Systems Biology. Academic Press, San Diego, pp. 211–228.

61. Pituello, F., Medevielle, F., Foulquier, F., Duprat, A.M., 1999. Activation of Pax6 depends on somitogenesis in the chick embryo cervical spinal cord. Development. 126, 587–596.

62. Royo, J.L., Maeso, I., Irimia, M., Gao, F., Peter, I.S., Lopes, C.S., D’Aniello, S., Casares, F., Davidson, E.H., Garcia-Fernandez, J., Gomez-Skarmeta, J.L., 2011. Transphyletic conservation of developmental regulatory state in animal evolution. PNAS. 108, 14186–14191.

63. Saad, R.S., Ghorab, Z., Khalifa, M.A., Xu, M., 2011. CDX2 as a marker for intestinal differentiation: Its utility and limitations. W. J. Gastroint. Sur. 3, 159–166.

64. Sakai, Y., Meno, C., Fujii, H., Nishino, J., Shiratori, H., Saijoh, Y., Rossant, J., Hamada, H., 2001. The retinoic acid-inactivating enzyme CYP26 is essential for establishing an uneven distribution of retinoic acid along the anterio-posterior axis within the mouse embryo. Genes Dev. 15, 213–225.

65. Sandmann, T., Girardot, C., Brehme, M., Tongprasit, W., Stolc, V., Furlong, E.E., 2007. A core transcriptional network for early mesoderm *Development* in Drosophila melanogaster. Genes Dev. 21, 436–449.

66. Sasai, N., Kutejova, E., Briscoe, J., 2014. Integration of signals along orthogonal axes of the vertebrate neural tube controls progenitor competence and increases cell diversity. PLoS Bio. 12, e1001907.

67. Savory, J.G., Bouchard, N., Pierre, V., Rijli, F.M., De Repentigny, Y., Kothary, R., Lohnes, D., 2009. Cdx2 regulation of posterior development through non-Hox targets. Development. 136, 4099–4110.

68. Scardigli, R., Baumer, N., Gruss, P., Guillemot, F., Le Roux, I., 2003. Direct and concentration-dependent regulation of the proneural gene Neurogenin2 by Pax6. Development. 130, 3269–3281.

69. Schneider, C.A., Rasband, W.S., Eliceiri, K.W., 2012. NIH Image to ImageJ: 25 years of image analysis. Nat. Meth. 9, 671–675.

70. Shimizu, T., Bae, Y.K., Hibi, M., 2006. Cdx-Hox code controls competence for responding to Fgfs and retinoic acid in zebrafish neural tissue. Development. 133, 4709–4719.

71. Shu, J., Fu, H., Qiu, G., Kaye, P., Ilyas, M., 2013. Segmenting overlapping cell nuclei in digital histopathology images. IEEE Eng. Med. Bio. Soc. Ann. Conf. 2013, 5445–5448.

72. Skaggs, K., Martin, D.M., Novitch, B.G., 2011. Regulation of spinal interneuron development by the Olig-related protein Bhlhb5 and Notch signaling. Development. 138, 3199–3211.

73. Skromne, I., Thorsen, D., Hale, M., Prince, V.E., Ho, R.K., 2007. Repression of the hindbrain developmental program by Cdx factors is required for the specification of the vertebrate spinal cord. Development. 134, 2147–2158.

74. Spann, P., Ginsburg, M., Rangini, Z., Fainsod, A., Eyal-Giladi, H., Gruenbaum, Y., 1994. The spatial and temporal dynamics of Sax1 (CHox3) homeobox gene expression in the chick’s spinal cord. Development. 120, 1817–1828.

75. Tamashiro, D.A., Alarcon, V.B., Marikawa, Y., 2012. Nkx1-2 is a transcriptional repressor and is essential for the activation of Brachyury in P19 mouse embryonal carcinoma cell. Differentiation. 83, 282–292.

76. Tzouanacou, E., Wegener, A., Wymeersch, F.J., Wilson, V., Nicolas, J.F., 2009. Redefining the progression of lineage segregations during mammalian embryogenesis by clonal analysis. Dev. Cell. 17, 365–376.

77. van de Ven, C., Bialecka, M., Neijts, R., Young, T., Rowland, J.E., Stringer, E.J., Van Rooijen, C., Meijlink, F., Novoa, A., Freund, J.N., Mallo, M., Beck, F., Deschamps, J., 2011. Concerted involvement of Cdx/Hox genes and Wnt signaling in morphogenesis of the caudal neural tube and cloacal derivatives from the posterior growth zone. Development. 138, 3451–3462.

78. van den Akker, E., Forlani, S., Chawengsaksophak, K., de Graaff, W., Beck, F., Meyer, B.I., Deschamps, J., 2002. Cdx1 and Cdx2 have overlapping functions in anteroposterior patterning and posterior axis elongation. Development. 129, 2181–2193.

79. van Rooijen, C., Simmini, S., Bialecka, M., Neijts, R., van de Ven, C., Beck, F., Deschamps, J., 2012. Evolutionarily conserved requirement of Cdx for post-occipital tissue emergence. Development. 139, 2576–2583.

80. Wang, Y., Yabuuchi, A., McKinney-Freeman, S., Ducharme, D.M., Ray, M.K., Chawengsaksophak, K., Archer, T.K., Daley, G.Q., 2008. Cdx gene deficiency compromises embryonic hematopoiesis in the mouse. PNAS. 105, 7756–7761.

81. White, R.J., Nie, Q., Lander, A.D., Schilling, T.F., 2007. Complex regulation of cyp26a1 creates a robust retinoic acid gradient in the zebrafish embryo. PLoS Bio. 5, e304.

82. Wilkinson, D.G., Nieto, M.A., 1993. Detection of messenger RNA by in situ hybridization to tissue sections and whole mounts. Meth. Enzy. 225, 361–373.

83. Wilson, V., Olivera-Martinez, I., Storey, K.G., 2009. Stem cells, signals and vertebrate body axis extension. Development. 136, 1591–1604.

84. Yamaguchi, T.P., Takada, S., Yoshikawa, Y., Wu, N., McMahon, A.P., 1999. T (Brachyury) is a direct target of Wnt3a during paraxial mesoderm specification. Genes Dev. 13, 3185–3190.

85. Young, T., Rowland, J.E., van de Ven, C., Bialecka, M., Novoa, A., Carapuco, M., van Nes, J., de Graaff, W., Duluc, I., Freund, J.N., Beck, F., Mallo, M., Deschamps, J., 2009. Cdx and Hox genes differentially regulate posterior axial growth in mammalian embryos. Dev. Cell. 17, 516–526.

## References

1. Bel-Vialar, S., Medevielle, F., Pituello, F., 2007. The on/off of Pax6 controls the tempo of neuronal differentiation in the developing spinal cord. Dev. Bio. 305, 659–673.

2. Martinez-Morales, P.L., Diez del Corral, R., Olivera-Martinez, I., Quiroga, A.C., Das, R.M., Barbas, J.A., Storey, K.G., Morales, A.V., 2011. FGF and retinoic acid activity gradients control the timing of neural crest cell emigration in the trunk. J. Cell Bio. 194, 489–503.

3. Matsuda, T., Cepko, C.L., 2004. Electroporation and RNA interference in the rodent retina in vivo and in vitro. PNAS. 101, 16–22.

4. Megason, S.G., McMahon, A.P., 2002. A mitogen gradient of dorsal midline Wnts organizes growth in the CNS. Development. 129, 2087–2098.

5. Paik, E.J., Mahony, S., White, R.M., Price, E.N., Dibiase, A., Dorjsuren, B., Mosimann, C., Davidson, A.J., Gifford, D., Zon, L.I., 2013. A cdx4-sall4 regulatory module controls the transition from mesoderm formation to embryonic hematopoiesis. Stem cell Rep. 1, 425–436.

6. Tamashiro, D.A., Alarcon, V.B., Marikawa, Y., 2012. Nkx1-2 is a transcriptional repressor and is essential for the activation of Brachyury in P19 mouse embryonal carcinoma cell. Differentiation. 83, 282–292.

